# Live-cell single-molecule tracking highlights requirements for stable Smc5/6 chromatin association *in vivo*

**DOI:** 10.1101/2020.06.19.148106

**Authors:** Thomas J. Etheridge, Desiree Villahermosa, Eduard Campillo-Funollet, Alex Herbert, Anja Irmisch, Adam T. Watson, Hung Q. Dang, Mark A. Osborne, Antony W. Oliver, Antony M. Carr, Johanne M. Murray

## Abstract

The essential Smc5/6 complex is required in response to replication stress and is best known for ensuring the fidelity of homologous recombination. Using single-molecule tracking in live fission yeast to investigate Smc5/6 chromatin association, we show that Smc5/6 is chromatin associated in unchallenged cells and this depends on the non-SMC protein Nse6. We define a minimum of two Nse6-dependent sub-pathways, one of which requires the BRCT-domain protein Brc1. Using defined mutants in genes encoding the core Smc5/6 complex subunits we show that the Nse3 double-stranded DNA binding activity and the arginine fingers of the two Smc5/6 ATPase binding sites are critical for chromatin association. Interestingly, disrupting the ssDNA binding activity at the hinge region does not prevent chromatin association but leads to elevated levels of gross chromosomal rearrangements during replication restart. This is consistent with a downstream function for ssDNA binding in regulating homologous recombination.

## Introduction

The structural maintenance of chromosomes (SMC) complexes cohesin, condensin and Smc5/6 are critical for the correct organisation of chromosome architecture^1^. Whereas the functions of cohesin and condensin are increasingly well understood, the exact function of Smc5/6 complex remains relatively ambiguous. Smc5/6 is conserved across all eukaryotes and is best known for its role in the cellular response to DNA damage by ensuring the fidelity of homologous recombination repair (HRR)^2,3^. Smc5/6 has been reported to promote replication fork stability^4^ and facilitate DNA replication through natural pausing sites^5^. Biochemically, the complex can regulate pro-recombinogenic helicases^6,7^. It has also been proposed to monitor DNA topology^8^ and recently been shown to restrict viral transcription^9,10^. Hypomorphic mutants show significant defects in sister-chromatid HRR, display replication fork instability, are sensitive to a wide range of genotoxins and accumulate unresolved recombination intermediates^4,11,12^. Intriguingly, complete inactivation of the Smc5/6 complex in a variety of organisms leads to cell death and this essential nature suggests it possesses additional functions beyond HR as deletions of core HR factors are viable.

Like all SMC complexes, the core of Smc5/6 is composed of two folded proteins, Smc5 and Smc6, which form a heterodimer (Figure 1A). Each subunit comprises a long coiled-coil arm with a hinge region at one end and a globular ATPase head at the other^1^. All three SMC heterodimers interact at the hinge and ATP binding/hydrolysis occurs in two pockets formed between the heads of the two subunits. For all SMC complexes, ATP turnover is essential for cell viability and has been proposed to bring about conformational changes in the arms^13–15^. The ATPase activity is also key to the interaction of SMC’s with DNA: cohesin’s ATPase is required for both loading and dissociation from DNA^16^, whilst condensin is dependent on its ATPase activity for translocating along DNA and forming loop structures^17,18^. The role of the Smc5/6 ATPase in DNA association has not been studied in detail.

**Figure 1.**
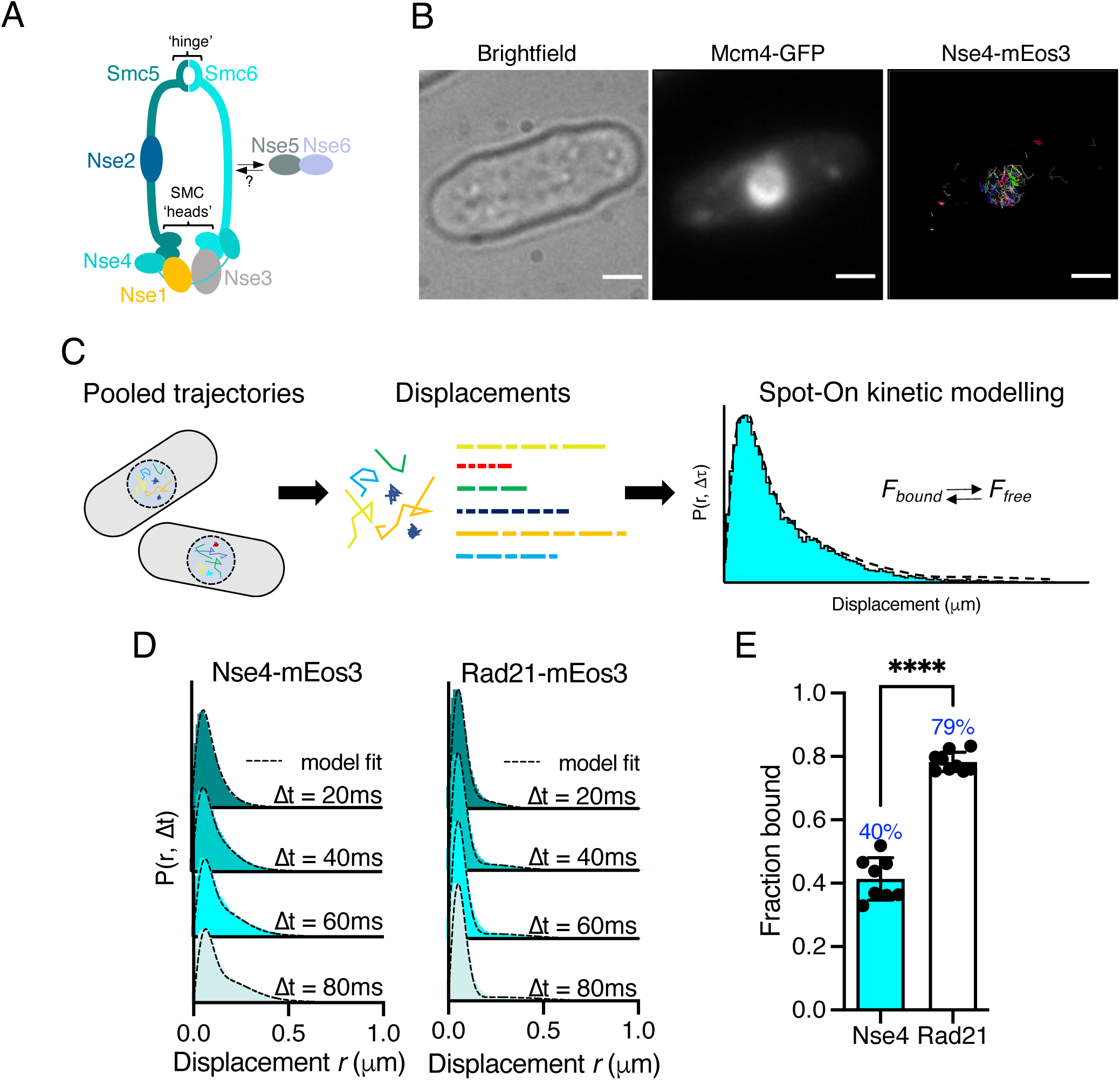
Single particle tracking of Smc5/6 to monitor chromatin association in live cells. A. Schematic representation of the Smc5/6 complex in fission yeast. B. Nse4-mEos3 tracking shows nuclear localisation of trajectories. SPT trajectories demonstrated confinement within nuclear region (right), that colocalised with the nuclear replication protein Mcm4 fused to GFP. Scale bar = 2μm C. Overview of approach to quantifying chromatin association using SPT data and Spot-On kinetic modelling. D. Probability density function (PDF) histograms and Spot-On model fitting (dashed line) for Nse4-mEos3 (Smc5/6) and Rad21-mEos3 (cohesin) single-molecule displacements at different time intervals. Displacements are from 3 pooled independent experiments, each with three technical repeats. E. Fraction bound values derived from Spot-On model fitting. Mean (+/− S.D). Black dots indicate Spot-On F_bound_ values derived from each technical repeat from 3 independent experiments. Percentages in blue denote fraction bound value from fitting pooled data in D. **** = p<0.0001.

The Smc5/6 hinge contains specialised interfaces that are important for interacting with single stranded DNA (ssDNA)^19^. Disruption of these regions by mutation results in sensitivity to DNA damaging agents. The Smc5/6 ATPase heads are bridged by a sub-complex of three <u>n</u>on-<u>S</u>MC <u>e</u>lements (Nse), Nse4 (kleisin) and two kleisin-interacting tandem winged-helix element (KITE) proteins, Nse1 and Nse3. Nse1 has a RING finger and, in association with Nse3, has been shown to have ubiquitin ligase activity^20^. The winged-helix domain of Nse3 possesses double-stranded DNA (dsDNA) binding activity, which is essential for viability^21^. The dsDNA binding has been predicted to provide the basis for initial chromatin association and loading of the complex^21^. In addition to the Nse1/3/4 subcomplex, Nse2, a SUMO ligase, is associated with the Smc5 coiled-coil arm. DNA association of the Smc5/6 complex is required to activate the Nse2 SUMO ligase, which SUMOylates a range of targets within and outside of the complex^22^. Two further proteins, Nse5 and Nse6, also associate with the Smc5/6 complex in yeasts (both *Saccharomyces cerevisiae* and *Schizosaccharomyces. pombe*). However, unlike the other Nse proteins, Nse5 and Nse6 have not been identified as part of a Smc5/6 holo-complex in human cells^23,24^.

Chromatin loading of the structurally related cohesin complex requires accessory proteins, the cohesin–loader complex Scc2–Scc4 (*sp*Mis4–Ssl3)^25^. A loading complex for Smc5/6 has not yet been defined but recent work in fission yeast has shown that its recruitment to sites of replication fork collapse occurs via a multi-BRCT domain protein, Brc1^26^. Brc1 binds to γ-H2A and interacts with the Nse5-Nse6 subcomplex (which associates with Smc5/6 but is not part of the core complex), providing a potential mechanism by which Smc5/6 is recruited and loaded. In *S. cerevisiae* the N-terminal four BRCT domains of the Brc1 homologue, Rtt107, have also been shown to bind Nse6 amongst a number of other proteins in the DNA damage response^27^. In human cells recruitment of Smc5/6 to inter-strand cross-links was shown to depend on interactions between SLF1, another multi-BRCT domain protein, and SLF2 - a distant homologue of Nse6^28^. These observations suggest that recruitment of Smc5/6 through Nse6 and a BRCT-domain mediator protein has been conserved through evolution.

Understanding how Smc5/6 is recruited to, and associates with, the chromatin is an important step in defining how it regulates recombination processes and other potential DNA transactions. To date, the study of Smc5/6 chromatin association has been mostly limited to chromatin immunoprecipitation (ChIP)-based methodologies. Recent studies have shown single-particle tracking (SPT) microscopy can provide robust measurements of chromatin interacting proteins in vivo and offer complementary data to genome-wide approaches.

Here, we perform SPT using photoactivated localisation microscopy (PALM) in live fission yeast cells to monitor chromatin association of Smc5/6. Using a range of *smc* and *nse* mutants we investigated the role of its ATPase activity, DNA interaction sites and protein binding partners in promoting chromatin association. This highlighted that ATPase activity and dsDNA binding are both crucial for chromatin association. In contrast, interaction with ssDNA at the hinge is not required for stable chromatin loading but we show that it is important to prevent gross chromosomal rearrangements at collapsed replication forks. We also establish that the Nse5-Nse6 sub-complex is required for almost all chromatin association, whereas Brc1 is required for only a proportion of the association. These data define the Brc1-Nse6-dependent sub-pathway of chromatin interaction and identify parallel Nse6-dependent but Brc1-independent sub-pathway(s).

## Results

### Smc5/6 is chromatin associated in unchallenged cells

To monitor Smc5/6 chromatin association in living yeast cells we used photoactivated localisation microscopy combined with single-particle tracking (SPT)^29^. We created a fission yeast strain that endogenously expressed the kleisin subunit Nse4 fused to the photoconvertible fluorophore mEos3 and verified this allele had no measurable impact on cellular proliferation (Figure S1A). We imaged photoconverted subsets of Nse4-mEos3 in live yeast cells at high temporal resolution (20ms exposure) and created trajectories by localising and tracking individual fluorophores (Figure S2A, B). Nse4-mEos3 localisations and trajectories showed nuclear confinement consistent with previous studies^30^ (Figure 1B).

To evaluate the chromatin association of Smc5/6 from our data we used the recently described ‘Spot-On’ software^31^ (see materials and methods). Spot-On implements a bias-aware kinetic modelling framework and robustly extracts diffusion constants and subpopulations from histograms of the molecular displacements that make up each trajectory (Figure 1C). We tracked Nse4-mEos3 in asynchronous live cells and created displacement histograms over 4-time intervals (Figure 1D). The profiles show a clear peak of short displacements (<100nm) indicative of a chromatin-bound fraction of Nse4-mEos3 in unchallenged cells. Spot-On kinetic modelling revealed a fraction bound of about 40% (Figure 1E). The displacement distributions were best described with a 3-state fit which, in addition to bound and freely diffusing species, included an intermediate slow-diffusing population. This may describe transient interactions with chromatin or anomalous diffusion as a result of a crowded molecular environment^32^ (Figure S8, materials and methods). Tracking of other core Smc5/6 (Nse2 and Smc6) subunits revealed similar displacement profiles and bound fractions, suggesting the dynamics of the kleisin subunit is indicative of the whole complex (Figure S3A).

We next compared Smc5/6 chromatin association to the structurally related cohesin complex. As fission yeast cells reside in G2 for the majority of the cell cycle we hypothesised that cohesin would be stably associated with the chromatin^33^ and should thus demonstrate a higher fraction bound. As predicted, tracking of Rad21 (kleisin) and Smc1 (arm) fused to mEos3 revealed displacement profiles with greater proportions of short displacements compared to Smc5/6 subunits and subsequently resulted in greater bound fractions extracted from Spot-On model fitting (Figure 1D and E, S3B). These observations show that interaction of cohesin and Smc5/6 with chromatin are distinct and different and suggest that their association occurs with different dynamics.

### dsDNA binding is required for efficient chromatin association

Smc5/6 has been shown to bind both ds- and ssDNA. The KITE protein Nse3 has a dsDNA binding domain in both humans and fission yeast and is situated at the head end of the complex (Figure 2A). This activity is essential and was predicted to be the initial point of interaction between Smc5/6 and the chromatin required before loading^21^. To assess whether Nse3 dsDNA interaction plays a role in global chromatin association we tracked Nse4-mEos3 in a *nse3-R254E* genetic background. This hypomorphic mutation has been shown to disrupt but not fully abolish dsDNA binding by Nse3^21^. When compared to *nse3*^+^, Nse4-mEos3 displacement histograms from asynchronous *nse4-mEos3 nse3-R254E* cells showed a broader profile suggesting the complex had become more dynamic (Figure 2B, C). This resulted in a reduction in the fraction bound value in Spot-On analysis (Figure 2D). This confirms *in vivo* that dsDNA binding by Nse3 underpins the chromatin association of Smc5/6.

**Figure 2.**
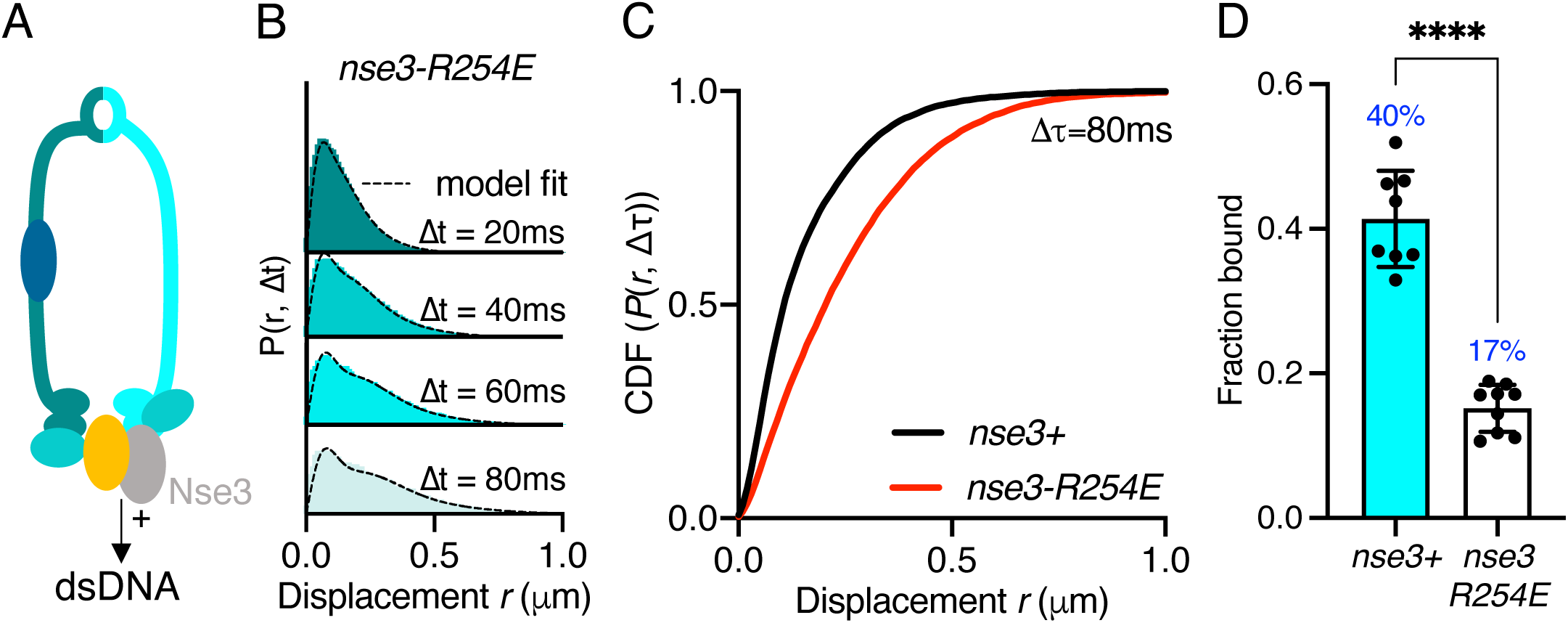
Stable Smc5/6 chromatin association requires dsDNA binding activity. A. Schematic representation of the region of known dsDNA interaction in *S. pombe* Smc5/6. B. Probability density function histogram of pooled Nse4-mEos3 single-molecule in *nse3-R254E* background and Spot-On model fitting (dashed line). the resulting fraction of bound molecules compared to wild type data set. Bar chart shows mean +/− S.E.M. Black dots denote independent repeats. *** p=0.0003 C. Cumulative distribution function (CDF) of pooled Δt = 80ms data from B. D. F_bound_ values derived from Spot-On model fitting of Nse4-mEos3 in *nse3-R254E* background. Black dots denote each technical repeat from 3 independent experiments. Percentages in blue denote fraction bound value from fitting pooled data in B. Mean (+/− S.D). **** = p<0.0001.

### Smc5/6 ATPase activity is required for efficient chromatin association

Each of the SMC complexes possess ATPase activity, with two separate and distinct active sites within juxtaposed ‘head’ domains, which are generated by bringing together the required signature motifs *in trans* (Figure 3A). Like all SMC complexes the ATPase activity of Smc5/6 is essential and inactivating mutations in either of the two Walker motifs are non-viable^34,35^. Therefore, to investigate the influence of ATPase activity on chromatin-association of the Smc5/6 complex, we first mutated the ‘arginine-finger’ of Smc5 (*smc5-R77A*) or Smc6 (*smc6-R150A*). Mutation of the equivalent residues in other SMC complexes does not typically affect the basal level of ATP turnover, but instead acts to abolish stimulation of activity by DNA-interaction^36^. Both the *smc5-R77A* and the *smc6-R150A* mutation resulted in sensitivity to replication stress (Figure S4A). Tracking of Nse4-mEos3 in these genetic backgrounds revealed increased single molecule displacements and subsequent decreases in chromatin association of the Smc5/6 complex (Figure 3B, C and S4B). *smc6-R150A* led to a dramatic decrease in chromatin association whereas mutation of the Smc5 arginine was noticeably less detrimental. Interestingly, the reduction in the levels of chromatin association correlated with sensitivity to exogenous genotoxic agents, strongly suggesting that DNA-dependent ATP hydrolysis by the two binding pockets is not equivalent.

**Figure 3.**
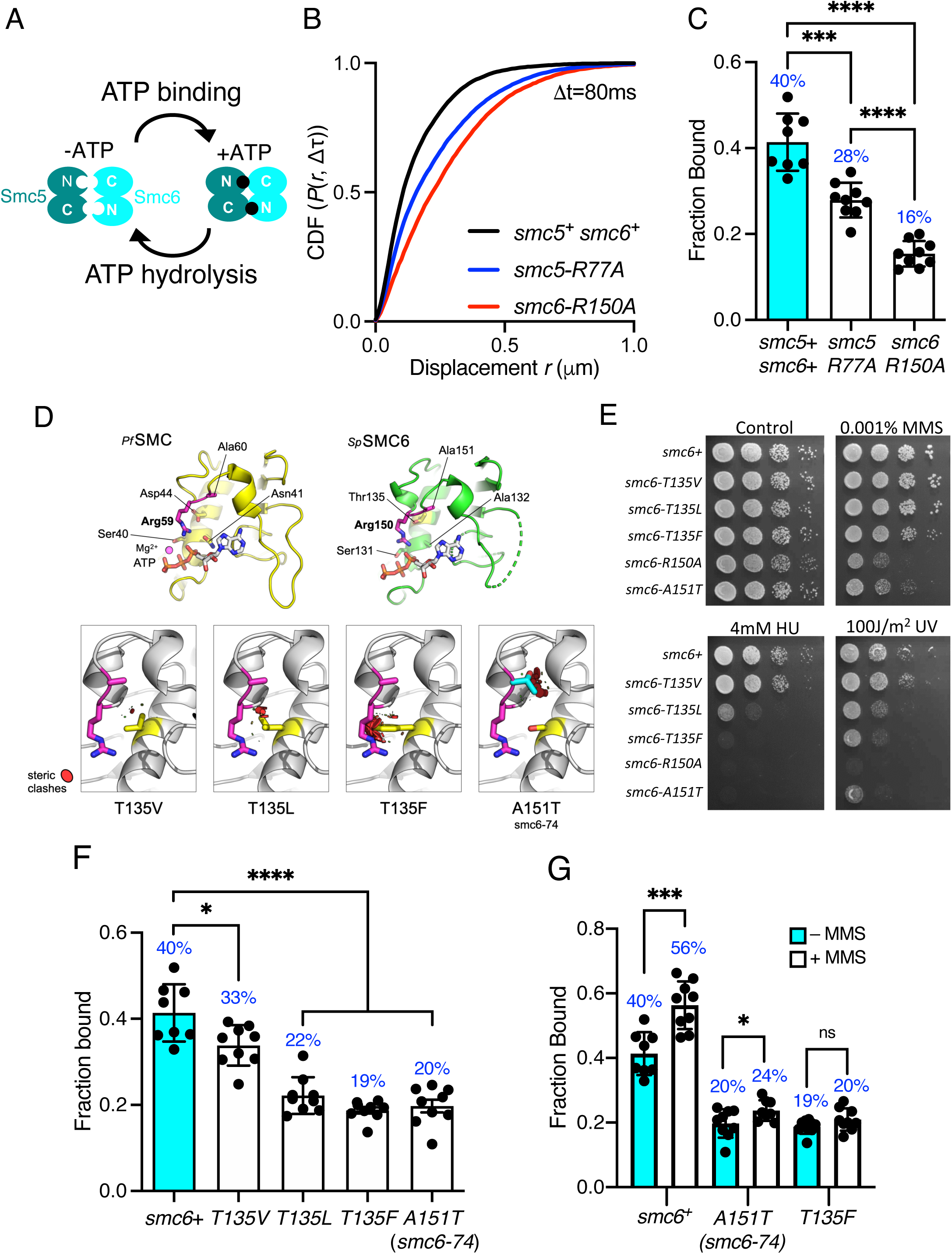
Smc5/6 ATPase activity regulates chromatin association. A. Schematic representation of SMC head engagement upon ATP binding. B. CDF of pooled Δt = 80ms single molecule displacements of Nse4-mEos3 in *smc5+ smc6+, smc6-R150A* and *smc5-R77A* genetic backgrounds. C. Comparison of the fraction of bound molecules from Nse4-mEos3 sptPALM experiments in asynchronous *smc6-R150A* and *smc5-R77A* genetic backgrounds to wild type dataset. Black dots denote each technical repeat from 3 independent experiments. Percentages in blue denote fraction bound value from fitting pooled data. Mean (+/− S.D). *** p=0.0001, **** = p<0.0001. D. Secondary structure molecular cartoons of homology models for the head domains of *S. pombe* Smc6, highlighting the arginine finger and its interaction with ATP. The X-ray crystal structure for the head domain of *Pyrococcus furiosus SMC* in complex with ATP served as a reference, providing the expected position of bound ATP the homology model. Key amino acids are shown in ‘stick representation’. The lower panel shows the predicted increase in severity of steric clashes made with the arginine finger through introduction of each of the indicated mutations. E. Yeast spot assay of *S. pombe* strains harbouring different *smc6* ATPase mutations grown at 30°C for 3 days. F. Fraction bound values in each of the *smc6*-*T135* mutant backgrounds compared to a wild type data set and *smc6-74* (A151T). Black dots denote each technical repeat from 3 independent experiments. Percentages in blue denote fraction bound value from fitting pooled data. Mean (+/− S.D). * p=0.0158, **** = p<0.0001. G. Fraction bound values derived from SPT analysis of MMS treated (0.03%, 5 hours) cells compared to asynchronous untreated data in F. * p=0.0495, *** p=0.0005

The Smc6 arginine finger mutant was of particular interest to us as the well characterised *smc6-74* allele maps to the next residue, A151T^4,34,38^. Single particle tracking showed this mutant to have a similar decrease in chromatin association to *smc6-R150A*. Sequence-threaded homology models for the head domain of *S. pombe* Smc6 and comparison to the X-ray crystal structure of the head domain from *Pyrococcus furiosus* SMC in complex with ATP (PfSMC, PDB: 1XEX) allowed us to create specific mutations designed to display a graduated effect on the Smc6 arginine-finger: Thr135 in Smc6 was mutated to a series of hydrophobic amino acids with increasing size, each predicted to produce increasingly severe steric clashes with the arginine-finger when engaged in interaction with bound ATP (Figure 3D).

Phenotypic analysis of each *smc6* mutant confirmed that the predicted severity of steric clash (Phe>Leu>Val) closely correlated with an increase in sensitivity to a range of genotoxic agents (Figure 3E), culminating with the most severe mutation, T135F, producing a phenotype similar to the well characterised *smc6-74* (A151T) mutant. Single-particle tracking data revealed that increasing the severity of the substitution corresponded with a decrease in the fraction of bound Smc5/6 (Figure 3F, S4C). The *smc6-T135F* strain showed similar levels of bound complex as the *smc6-74* mutation.

Since mutations in the ATPase domains render cells sensitive to replication stress (Figure 3E) we monitored whether these mutants could recruit the complex to chromatin after treatment with MMS. Acute exposure to 0.03% MMS for 5 hours resulted in a modest increase in the fraction of Nse4-mEos3 bound to the chromatin in cells with a wild type background (Figure 3G). However, in contrast both the *smc6-74* and *smc6-T135F* alleles significantly reduced or prevented Smc5/6 from being recruited to chromatin in response to MMS.

Together these data demonstrate that the ability to stimulate Smc5/6 ATPase activity through the arginine finger is crucial for its stable association with the chromatin. The disparity in phenotype between *smc6* and *smc5* ATPase mutants suggests there could be an underlying asymmetry in the use for the two ATP binding sites, a phenomenon that has been recently described for both condensin and cohesin^16,39^.

### ssDNA binding is dispensable for Smc5/6 chromatin association

We recently determined the structure of the *S. pombe* Smc5/6 hinge and demonstrated its preferential binding to ssDNA^19^. Specialised features known as the ‘latch’ and ‘hub’ are required for efficient association with ssDNA (Figure 4A). The kinetics of this interaction are biphasic and appear to involve two distinct interaction points. Like mutants compromised for dsDNA binding, mutations in these key regions that weaken the interaction with ssDNA render cells viable but sensitive to replication stress and DNA damaging agents^19^. We tested whether the ability to interact with ssDNA affected the ability of Smc5/6 to associate with chromatin.

**Figure 4.**
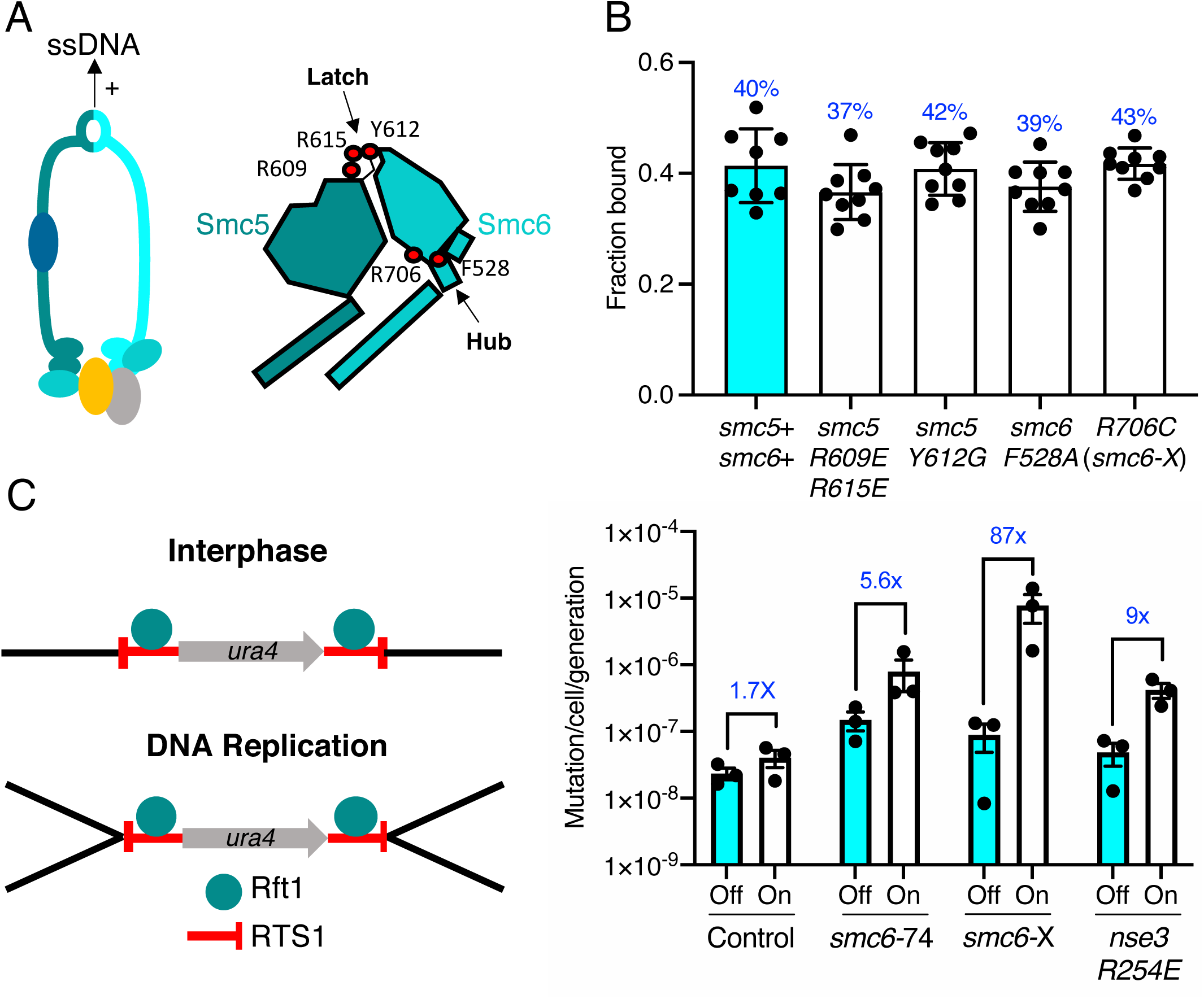
ssDNA interactions are required to prevent gross chromosomal re-arrangements but dispensable for stable Smc5/6 chromatin association. A. Left: Schematic representation of the hinge region known to interact with ssDNA interaction in Smc5/6. Right: Schematic diagram of the *S. pombe* hinge region adapted from^19^. Residues implicated in ssDNA interaction are highlighted with red filled circles. B. Fraction bound values of Nse4-mEos3 derived from SPT experiments in Smc5/6 hinge mutant backgrounds compared to wild type dataset. Mean +/− S.D. Black dots denote independent repeats and percentages in blue denote fraction bound value from fitting pooled data from all repeats. C. Diagram of the site-specific replication stall system *RTS1-ura4-RTS1*^40^, which consists of two inverted *RTS1* sequences integrated on either sides of the *ura4* gene. Rtf1 binds the *RTS1* sequence and stalls incoming replication forks coming from both centromeric and telomeric sides. Rtf1 is expressed under the control of the *nmt41* promoter which is “off” in the presence of thiamine and “on” upon thiamine removal. D. Induction of *rtf1* in cells harbouring *RuraR* construct induces *ura4* marker loss as assayed by 5-FOA resistance. Cells growing in the presence, (Off, arrest repressed) or absence (On, arrest induced) of thiamine were analysed by fluctuation analysis. Mean +/− S.E.M. Black dots denote independent repeats.

Previously characterised mutations were introduced into the Nse4-mEos3 strain that affect either initial ssDNA interaction (*smc5-R609E R615E*), stable hinge heterodimerisation (*smc5-Y612G*) or secondary ssDNA interactions at the Smc6 hub (*smc6-F528A, smc6-R706C*)^19^ (Figure 4A, right). Spot-On model fitting to sptPALM data showed that, unlike the dsDNA binding and ATPase mutants, disruption of ssDNA interactions did not alter the bound fraction of Smc5/6 in unchallenged cells (Figure 4B).

Since these mutations render cells sensitive to replication stress, we monitored recruitment of Smc5/6 complex to chromatin after treatment with MMS. Disruption of ssDNA interactions either reduced, or prevented, further Smc5/6 from being recruited to chromatin in response to MMS (Figure S5). Together, these data show that, while dsDNA binding is required for stable association of the Smc5/6 complex with chromatin, its interactions with ssDNA are not. This is consistent with ssDNA interactions playing a role in processes downstream of loading and we speculate that it may be important for Smc5/6 retention on the DNA during DNA repair-associated processes.

### ssDNA interaction is required to prevent gross chromosomal rearrangements

We hypothesised that the loss of Smc5/6 chromatin association would produce distinct outcomes during HR-dependent processes compared to the loss of ssDNA interaction. To investigate this, we compared the effect of Smc5/6 mutations in the response to replication fork stalling in the previously characterised *‘RuraR’* replication fork barrier system^40^.

In fission yeast, binding of Rtf1 to the replication termination sequence, *RTS1*, arrests replication forks in a polar manner^41^. In the *RuraR* system, two copies of *RTS1* are placed in an inverted orientation on either side of the *ura4* marker on chromosome III (Figure 4C). The *RTS1* barrier activity is regulated by placing *rtf1* under the control of the *nmt41* promoter and induction of *rtf1* expression leads to arrest of replication forks converging on both *RTS1* sequences. Replication of the intervening *ura4* requires homologous recombination-dependent replication restart which can result in genome instability via non-allelic homologous recombination (NAHR)^42^ or small scale errors by the error prone restarted fork^43^. The loss of *ura4* in the *RuraR* system provides a readout that is particularly useful to characterise NAHR events. In the absence of key HR factors, such as Rad51, induction of arrest leads to viability loss, whereas mis-regulation of HR generates aberrant outcomes^40^.

We introduced the *smc6-R706C* (*smc6-X*) and *smc6-A151T* (*smc6-74*) mutations into the *RuraR* system. There was no loss of viability when stalling was induced at *RTS1* in these backgrounds compared to *rad51*Δ (Figure S6A). This is consistent with Smc5/6 regulating recombination, rather than being core to the recombination process^2^. Induction of replication arrest led to an increase in the loss of *ura4* activity in *smc6*^+^, *smc6*-74 and *smc6*-X backgrounds. There was only a modest change in the ATPase mutant (*smc6*-74) (5.6-fold) compared to *smc6+* (1.7-fold) suggesting that reduced chromatin association only moderately effects HR fork restart. To confirm this further we introduced the *nse3-R254E* mutation into the *RuraR* strain and found similar results (9-fold) (Figure 4C).

Introduction of the hinge mutant (*smc6*-X) resulted in a highly elevated induction of *ura4* loss, an 87-fold increase over the uninduced (Figure 4C and Table S1). Analysis of *ura4*^-^ colonies isolated after replication stalling from *smc6-X* and *smc6-74* mutants showed that most were full deletions of the intervening sequence between the two *RTS1* loci (Figure S6 B and C). Thus, these results highlight the ssDNA binding region of the Smc5/6 hinge as particularly important for the suppression of NAHR and gross chromosomal rearrangements, and that stable recruitment of a defective complex (*smc6-*X) is more detrimental at collapsed replication forks than reduced Smc5/6 chromatin association (*smc6*-74 and *nse3-R254E*).

### Different requirements for Nse6 and Brc1 for recruitment of Smc5/6

Recent work in fission yeast has shown that the Nse6 subunit and the BRCT-containing protein Brc1 are required for the recruitment of Smc5/6 to distinct nuclear foci in response to DNA damage^26^ (Figure 5A). To investigate if these factors influence recruitment of the Smc5/6 complex to chromatin in unchallenged cells the genes encoding Brc1 and Nse6 were deleted in the Nse4-mEos3 strain and Smc5/6 chromatin association monitored by SPT.

**Figure 5.**
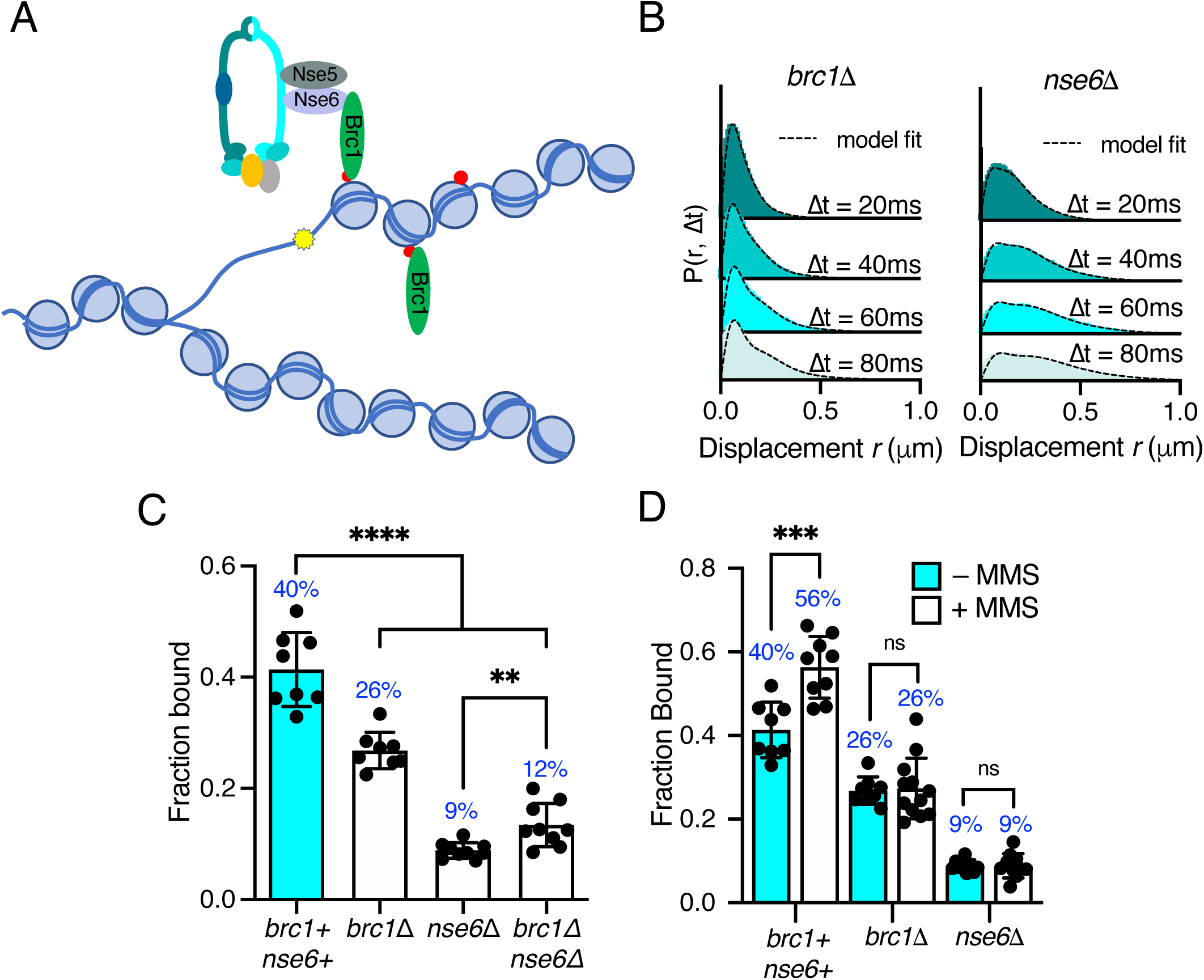
Differential requirements of Nse6 and Brc1 for Smc5/6 chromatin association. A. Schematic diagram of Smc5/6 recruitment to γ-H2A (red dots: H2A phosphorylation) at stalled replication forks. Brc1 binds to γ-H2A and recruits Smc5/6 via an interaction with Nse6. Yellow star indicates a DNA lesion. B. Displacement PDF histograms from asynchronous cells expressing Nse4-mEos3 in *brc1*Δ and *nse6*Δ genetic backgrounds. Data are from 3 pooled independent experiments, each with three technical repeats. Spot-On model fit is denoted by dashed line. C. Comparison of Nse4-mEos3 F_bound_ values derived from Spot-On fitting of SPT displacement histograms in wild type, *brc1*Δ*, nse6*Δ *and brc1*Δ *nse6*Δ genetic backgrounds. Mean +/− S.D. Black dots values derived from independent technical repeats, percentages in blue denote fraction bound value from fitting pooled data from all repeats. **** = p<0.0001, ** p=0.0043 D. F_Bound_ fraction values from *brc1*Δ *and nse6*Δ cells in C compared to parallel experiments where cells were treated with 0.03% MMS for 5 hours. *** = p<0.005, *ns* = not significant.

Deletion of either *brc1* or *nse6* resulted in an altered displacement profile and a concurrent decrease in the fraction of bound molecules (Figure 5B, C). In *brc1*Δ the amount of chromatin associated Smc5/6 decreased by approximately 35% showing that only a proportion of Smc5/6 chromatin association is dependent on Brc1. Recruitment of Brc1 to chromatin is reported to be via a specific interaction with γ-H2A^44^. We therefore investigated Smc5/6 complex recruitment in the absence of H2A phosphorylation. Introduction of *nse4-mEos3* into *hta1-SA hta2-SA* mutant cells revealed a statistically significant reduction in the fraction bound, similar to that seen in *brc1*Δ cells (Figure S7). These data are consistent with Brc1-dependant loading of Smc5/6 being largely confined to regions of γ-H2A.

In contrast, deletion of *nse6* showed significant deviation from the wild type data, resulting in an almost complete loss of chromatin associated Nse4 (Figure 5C), strongly supporting a Brc1-independent role for Nse6 in the stable recruitment of Smc5/6 to the chromatin. It should be emphasised that *nse6* deleted *S. pombe* cells are slow growing and very sensitive to genotoxins, whereas deletion of genes encoding proteins in the core complex are inviable. Deletion of *brc1* in an *nse6*Δ background is viable and results in additive sensitivity to DNA damage and replication stress^26^. This suggests that Smc5/6 can still associate with chromatin in the absence of Nse6, albeit at a severely reduced level. We hypothesise that the dsDNA binding activity of Nse3 is sufficient for this residual association with the chromatin. In support of this prediction, we were unable to generate the *nse6*Δ *nse3-R254E* double mutant suggesting it is synthetically lethal. Furthermore, SPT analysis of Nse4-mEos3 in *nse6*Δ *brc1*Δ cells did not lead to further reduction in the fraction of bound complexes (Figure 5D).

Previous ChIP experiments have shown that Smc5/6 is enriched at repetitive genomic loci following MMS treatment and that this is dependent on Brc1 and Nse6^26^. We tested whether we could detect increased Nse4 chromatin association in response to MMS treatment in *brc1*Δ and *nse6*Δ cells. Both *brc1*Δ and *nse6*Δ cells failed to show any increase above levels detected in untreated cells upon acute exposure to MMS (Figure 5D) supporting the hypothesis that both Brc1 and Nse6 are required for Smc5/6 recruitment to sites of DNA damage^26^.

### The Nse5-Nse6 subcomplex displays different kinetics than the Smc5/6 core complex

Intrigued by the significant role of Nse6 even in the absence of DNA damage we investigated the dynamics of the Nse5-Nse6 complex. We tagged both Nse5 and Nse6 with mEos3 (Figure S1B) and compared their behaviour to Nse4. In contrast to Nse2 and Smc6, which show similar chromatin association to Nse4 (Figure S3A), both Nse5 and Nse6 displayed a broader range of displacements and were subsequently less chromatin associated (Figure 6A and B). This suggests Nse6 is more dynamic than other subunits and may indicate its association with the core Smc5/6 complex is transient. To determine whether chromatin association of Nse5-Nse6 is affected by that of the core complex we introduced the *nse6-mEos3* allele into a *smc6-74* or *smc6-X* genetic background. We predicted that if Nse5-Nse6 was tightly associated with the core complex then it would display reduced association in a *smc6-74* strain as seen with Nse4, but not in *smc6-X* (Figure 3E and 4B). Tracking of Nse6-mEos3 in both mutants revealed no significant change in the fraction bound (Figure 6C) suggesting Nse5-Nse6 has different chromatin association dynamics to the core Smc5/6 complex. This would be indicative of Nse5-Nse6 acting to transiently stabilise or load Smc5/6 complexes on the chromatin.

**Figure 6.**
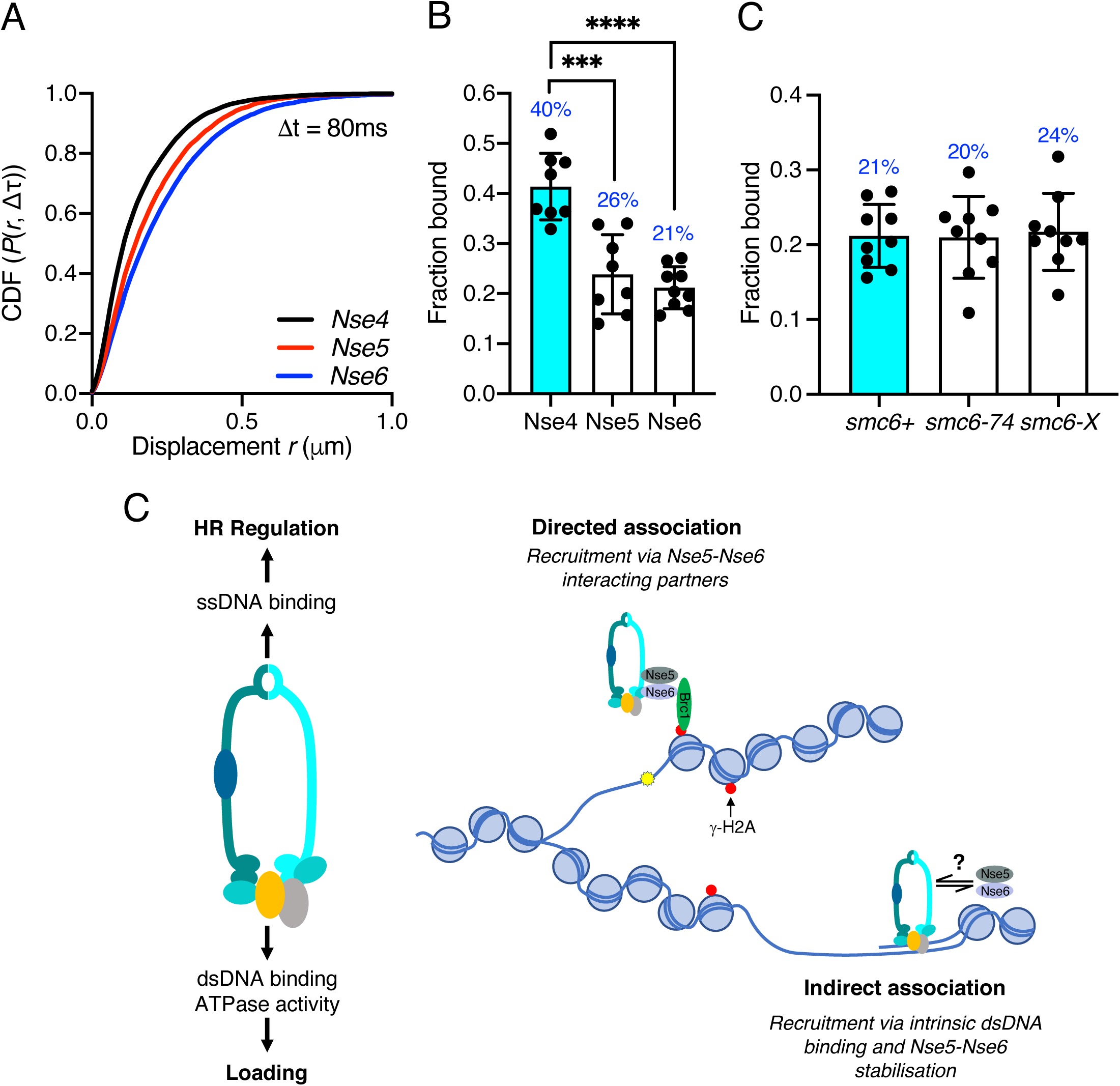
Nse5-Nse6 chromatin association is distinct from other Smc5/6 subunits. A. CDF histogram of pooled single molecule displacements at Δt = 80ms time interval of Nse4-mEos3, Nse5-mEos3 and Nse6-mEos3. B. Fraction of bound molecules extracted from Spot-On model fits from experiment in A. Mean +/− S.D. Black dots denote independent technical repeats, percentages denote fraction bound value from fitting pooled data from all repeats. *** p=0.0003, **** = p<0.0001 C. Fraction of bound molecules extracted from Spot-On model fits from SPT Nse6-mEos3 in *smc6-*74 or *smc6-*X genetic backgrounds compared to wild type data in B. D. Schematic diagram of Smc5/6 DNA interactions and their roles (left) and proposed model of Smc5/6 chromatin association (right). Loading requires dsDNA binding by Nse3 and Smc5 and Smc6 ATPase activity. ssDNA binding at the hinge is not required for loading but is required for subsequent functions to regulate homologous recombination, suppress non-allelic recombination and GCRs. Smc5/6 association with chromatin is dependent on Nse5 and Nse6 and either directed (e.g. Brc1-dependent recruitment to γ-H2A) (top) or non-directed via dsDNA binding and subsequent loading (bottom). Nse5/6 is required in both instances and may act either to directly load Smc5/6 or may stabilise its association after initial loading by dsDNA interaction.

## Discussion

The Smc5/6 complex is best known as a component of the DNA repair machinery that ensures the fidelity of homologous recombination (HR). However, the complex is essential in yeast which suggests it possesses additional functions beyond HR as deletions of core HR factors are viable^3^. The recruitment of Smc5/6 to DNA and ATP binding/hydrolysis at both the ATP sites are thought to be essential for each of its cellular roles. Understanding the molecular details of how Smc5/6 associates with DNA and/or chromatin is therefore an important step in elucidating how Smc5/6 regulates recombination and other potential DNA transactions. Here, we have established single-particle tracking as a method to probe Smc5/6 dynamics in live cells and coupled with yeast genetics and structural studies we uncover the key requirements for its association with chromatin.

### Smc5/6 complex features required for stable chromatin association

The Smc5/6 complex contains two separate ATP binding and hydrolysis sites. Both are formed when the Smc5 and Smc6 head domains interact. In common with all SMC complexes, the ATP binding pockets have an arginine finger, which is proposed to regulate DNA-dependent ATP hydrolysis. We show that mutating either of the Smc5 or Smc6 arginine fingers resulted in an increase in sensitivity to DNA damage and replication stress. This correlated with decreases in the fraction of bound Smc5/6 detected in SPT experiments. Interestingly, Smc5 and Smc6 arginine fingers were not equivalent as we uncovered an underlying asymmetry in the requirement of the two ATP binding sites for stable chromatin association. This asymmetry is in line with observations made for cohesin and condensin^18,39^.

One of the original *smc6* mutants, *smc6-74* (A151T) maps to the residue adjacent to the arginine residue in the arginine finger domain, suggesting it is compromised in ATP hydrolysis. Using a structural model based on the *Pyrococcus furiosus* SMC head domain, we engineered a series of structurally informed mutations designed to compromise the arginine finger to various degrees. This allowed us to dial in sensitivity to DNA damaging agents that robustly correlated with a reduced ability of Smc5/6 to associate with chromatin. Taken together, these observations strongly suggest that ATPase activity stimulated by DNA binding is pre-requisite for Smc5/6 complex DNA/chromatin association and function.

Recent structural and biophysical data for the ssDNA-binding activity of the Smc5/6 hinge domain^19^ and the dsDNA-binding Nse1/3/4 module^21^ allowed an investigation of the role for each of these two functions in promoting Smc5/6 chromatin association. The introduction of defined mutations into fission yeast demonstrated that dsDNA-binding by Nse3 is required for DNA/chromatin association of the Smc5/6 complex, whereas the ability to bind ssDNA at the hinge is dispensable. Since ssDNA-binding mutants are sensitive to a range of genotoxic agents^19^, we therefore predicted that ssDNA binding most likely plays a role in downstream processes once the complex has initially bound to dsDNA/chromatin. This would be an analogous situation to cohesin, whereby after initial DNA binding to dsDNA, capture of a second DNA moiety is only achievable for ssDNA^45^. This prediction is supported by results from our site-specific replication stall experiments, which indicate that increased levels of ectopic recombination occur in Smc5/6 mutants that lack the ability to interact with ssDNA correctly. This is much higher than in mutants that fail to stimulate ATPase activity and do not correctly associate with chromatin.

### Interacting factors influencing Smc5/6 chromatin association

Both Brc1 and Nse6 have been implicated in recruiting Smc5/6 to regions of γ-H2A at stalled/collapsed replication forks in fission yeast^26^. We demonstrate here that deletion of either one of these factors reduces the *in vivo* levels of chromatin-associated Smc5/6, in both unchallenged cells and after exposure to MMS. Interestingly, deletion of *brc1* or preventing histone H2A phosphorylation did not generate as severe a defect in chromatin association as deletion of *nse6*. This is in agreement with recent ChIP experiments performed at discreet genomic loci^26^ and demonstrates that there is at least one alternative Brc1-independent pathway for recruitment of Smc5/6 to chromatin.

To explain the data, we consider two possible modes of chromatin association: directed and non-directed association (Figure 6C). Directed association occurs when the complex is recruited to discrete genomic loci via interaction between the Nse5/6 subcomplex and chromatin associated factors. This occurs via Brc1 at sites of γ-H2A but alternative Nse5/6-interacting partners may exist to bring the complex to specific DNA structures, including stalled replication forks, HR intermediates and double strand breaks.

Association with the chromatin may also occur in a non-directed manner via Smc5/6’s intrinsic ability to associate with DNA through the dsDNA binding site of Nse3. In this scenario, Smc5/6 initially binds DNA structures directly via Nse3 and the Nse5/Nse6 subcomplex acts transiently to stabilise this interaction. This would help explain some important observations. Firstly, while Smc5, Smc6 and Nse1-4 are all essential proteins, fission yeast cells can survive without Nse5/Nse6. In the absence of Nse5/Nse6, the complex still possesses dsDNA binding activity, but the association with the chromatin is unstable. Secondly, deletion of *nse6* is synthetically lethal with the hypomorphic dsDNA binding mutant *nse3-R254E,* suggesting the dsDNA binding activity is sufficient to retain viability in the absence of exogenous DNA damage or replicative stress. If Nse5/6 is required to stabilise DNA/chromatin association after an initial recruitment by dsDNA binding, it would explain both the essential nature of the dsDNA binding activity of Nse3 and the observations that dsDNA binding site is tightly linked to chromatin association.

These two modes are not mutually exclusive and, in both cases, the Nse5/6 heterodimer may be acting transiently to regulate structural configurations of the complex that promote stable association with the chromatin (‘loading’), much like the model for Mis4-Ssl3 being the loader for cohesin^25,46^. Our SPT experiments show that Nse5 and Nse6 are more mobile than components of the core Smc5/6 complex suggesting alternative kinetics. This would be analogous to the cohesin loader Scc2 which displays different dynamics to the cohesin complex and ‘hops’ between chromatin bound cohesin molecules^47^. Intriguingly, two recent studies have demonstrated that Nse5/6 negatively regulates the ATPase activity of Smc5/6 *in vitro*, and binding to the core complex causes conformational alterations^48,49^. Taken together with our observations that DNA-stimulated ATPase activity is required for stable loading to the chromatin, this provides an Nse5/6-dependent mechanism by which ATPase activity is repressed until a DNA substrate is encountered. We predict that once Nse5/6 inhibition of Smc5/6 ATPase is relieved it is then released from the core complex.

In summary, by conducting a detailed characterisation of Smc5/6 chromatin association in live cells we demonstrate that SPT is a powerful approach for studying this enigmatic complex. This methodology, when coupled with structure-led mutational analysis and yeast genetics, has provided new insights into Smc5/6 behaviour as well as clarifying previous observations from past genetic and molecular genetic experiments.

## Materials and Methods

### S. pombe strain construction

*S. pombe* strains were constructed using Cre-lox mediated cassette exchange (RMCE) as previously described^50^. Strains were created either with essential gene replacement base strains or C-terminal tagging base strains (Supplementary table S2). C-terminal base strains were transformed with plasmid pAW8-mEos3.2-KanMX6 to introduce the mEos3.2 tag at the C-terminal end of the gene.

### Microscopy sample preparation

*S. pombe* cultures were grown to mid-log phase at 30°C in Edinburgh minimal media (EMM) supplemented with leucine, uracil and adenine. Cells were harvested and washed once in phosphate buffered saline (PBS). Cells were then resuspended in PBS and 10μl was deposited on an EMM-agarose pad before being mounted on ozone-cleaned circular coverslips (Thorlabs, #1.5H, ∅25mm) and placed in a metal cell chamber for imaging (Attofluor, Thermofisher). For replicative stress experiments, MMS was added to cultures at a final concentration of 0.03% and incubated for 5 hours before being processed for imaging.

### PALM microscopy

Live *S. pombe* cells were imaged with a custom-built microscope similar to that previously described^51^. The microscope is built around an inverted Olympus IX73 body fitted with a motorized stage (Prior H117E1I4) and a heated incubation chamber (Digital Pixel Ltd). Cells were illuminated using a 561-nm imaging laser (Cobolt, Jive) and a 405-nm activation laser (LaserBoxx, Oxxius). Both laser beams were expanded and collimated and were focused to the back focal plane (BFP) of an apochromatic 1.45 NA, 60× TIRF objective (Olympus, UIS2 APON 60× OTIRF). Both beams were angled in a highly inclined near-TIRF manner to achieve high signal-to-background. Illumination of the sample was controlled via mechanical shutters and all components were computer-controlled using the Micro-Manager software. The emission fluorescence from the sample was filtered with a band-pass filter (Semrock 593/40) before being expanded to create an optimized image pixel size of 101 nm after projection onto the EMCCD camera (Photometrics Evolve 512 Delta).

Samples were mounted on microscope stage and incubated at 30°C. Cells were illuminated with continuous 561nm excitation (8.3mW at rear aperture of objective lens) and pulsed with 100ms 405nm laser illumination every 10s in order to photoconvert mEos3.2 molecules (max. 0.23mW at rear aperture of objective lens). We established the number of nuclei that needed to be assayed for reproducibility empirically. To ensure that single-molecule traces were recorded from a sufficient number of nuclei (>50) each biological repeat consisted of data collection from at least 2 separate fields of view imaged one after the other (technical repeats). Each acquisition consisted of 20,000 frames with a camera exposure time of 20ms.

### Single particle tracking data analysis

Raw sptPALM data was analysed using the ‘PeakFit’ plugin of the GDSC single-molecule localisation microscopy software package for Fiji (GDSC SMLM - https://github.com/aherbert/gdsc-smlm). Single molecules were identified and localised using a 2D gaussian fitting routine (configuration file available on request). Nuclear localisations consisting of a minimum of 20 photons and localised to a precision of 40nm or better were retained for further analysis. Single molecules were then tracked through time using the ‘Trace Diffusion’ GDSC SMLM plugin. Localisations appearing in consecutive frames within a threshold distance of 800nm were joined together into a trajectory^51^. Single molecule trajectories were then exported into .csv Spot-On format using the ‘Trace Exporter’ plugin.

Track data was uploaded into the Spot-On web interface and was analysed using the following jump length distribution parameters: Bin width (μm) =0.01, number of timepoints =5, Jumps to consider =4, Max jump (μm) =3. For all Smc5/6 components, data sets were fit with a 3-state Spot-On model using the default parameters, except for: D_slow_ min =0.08, localisation error fit from data =yes, dZ (μm) =0.9. The decision on which Spot-On model to fit was based on the Akaike information criterion (AIC) reported by Spot-On (see Figure S8). It is not clear whether this third state describes transient interactions with chromatin or arises from anomalous diffusion as a result of a crowded molecular environment^32^. For cohesin data sets we fit a two-state model with the same parameters, excluding D_slow_. In all cases, the model was fit to the cumulative distribution function (CDF).

Probability density function (PDF) histograms and model fit were created using data combined from all three repeats of an experiment and exported from Spot-On before being graphed in Prism (GraphPad). Bar charts were produced by fitting data collected in each repeat (three fields of view) and extracting the fraction of bound molecules. Black circles represent the value derived for each repeat, bars represent the mean and error bars denote standard error of the mean. Two-tailed t-test was performed in Prism software of the Spot-On F_bound_ values from three repeats. Nuclear single molecule traces used for analysis in SpotOn are available via the Open Science Framework (osf.io/myxtr).

### Structural modelling

Sequence-threaded homology models for the head domains of both *S. pombe* Smc5 and Smc6 were generated using the PHYRE2 web portal^52^. The potential effects of introducing single point mutations were assessed using PyMOL (v2.32, The PyMOL Molecular Graphics System, Version 2.32, Schrödinger, LLC)

### Yeast spot test assay

Yeast strains were cultured in yeast extract (YE) overnight to mid-log phase. Cells were harvested and resuspended to a concentration of 10^7^ cells/ml. Serial dilutions were then spotted onto YE agar plates containing the indicated genotoxic agent.

### Yeast gross chromosomal rearrangement assay

The rate of *ura4^+^* loss in the *RuraR* system was measured using a previously described fluctuation test^40^. Colonies growing on YNBA plates lacking uracil (and containing thiamine) were re-streaked onto YNBA plates containing uracil, either in the presence or absence of thiamine. After 5 days, 5 colonies were picked from either condition and each was grown to saturation (∼48hrs) in 10ml liquid EMM culture containing uracil, with or without thiamine.

Each culture was counted and about 1×10^7^ cells were plated in triplicate on YEA plates containing 5’-fluoroorotic acid (5’-FOA; Melford). 100μl of a 1:20000 dilution of each saturated culture (about 200 cells) was plated in duplicate on YEA as titre plates. After 5 to 7 days of growth, 5-FOA resistant colonies and colonies on YEA were counted. A proportion of 5-FOA resistant colonies were streaked on YNBA lacking uracil to verify *ura4* gene function loss. These *ura4^-^* colonies were used in the translocation PCR assay as described previously^40^. The rate of *ura4* loss per cell per generation was calculated using the maximum likelihood estimate of the Luria-Delbruck with a correction for inefficient plating^53^. We performed all computations using the R package rSalvador^54^.

## Author contributions

TJE, AWO, AMC and JMM conceived the experimental approach. TJE and MAO built the custom microscope. TJE acquired and analysed the microscopy data. AH wrote and benchmarked the PeakFit custom single-molecule software. TJE, DVC, AI, HQD and ATW performed strain construction, phenotypic analyses and molecular biology. AI performed translocation PCR assay. ECF performed fluctuation test data analysis. AWO performed structural analysis and designed mutations. TJE, AWO, AMC and JMM wrote the manuscript.

## Acknowledgements

We would like to acknowledge Anders Hansen and Maxime Woringer for their help with initial implementation of the Spot-On software for analysis of SPT data in fission yeast. We thank Sarah Lambert for control strains for the GCR assay and Steven F. Lee for single-molecule microscopy advice. We also thank J. Palecek for the *nse3-R254E* strain. AMC acknowledges support from the Wellcome trust (110047/Z/15/Z), JMM and AWO acknowledge support from the MRC (MR/P018955/1). TJE would like to dedicate this manuscript to the memory of Vivien Horobin.

---

**Supplementary Figure 1.**
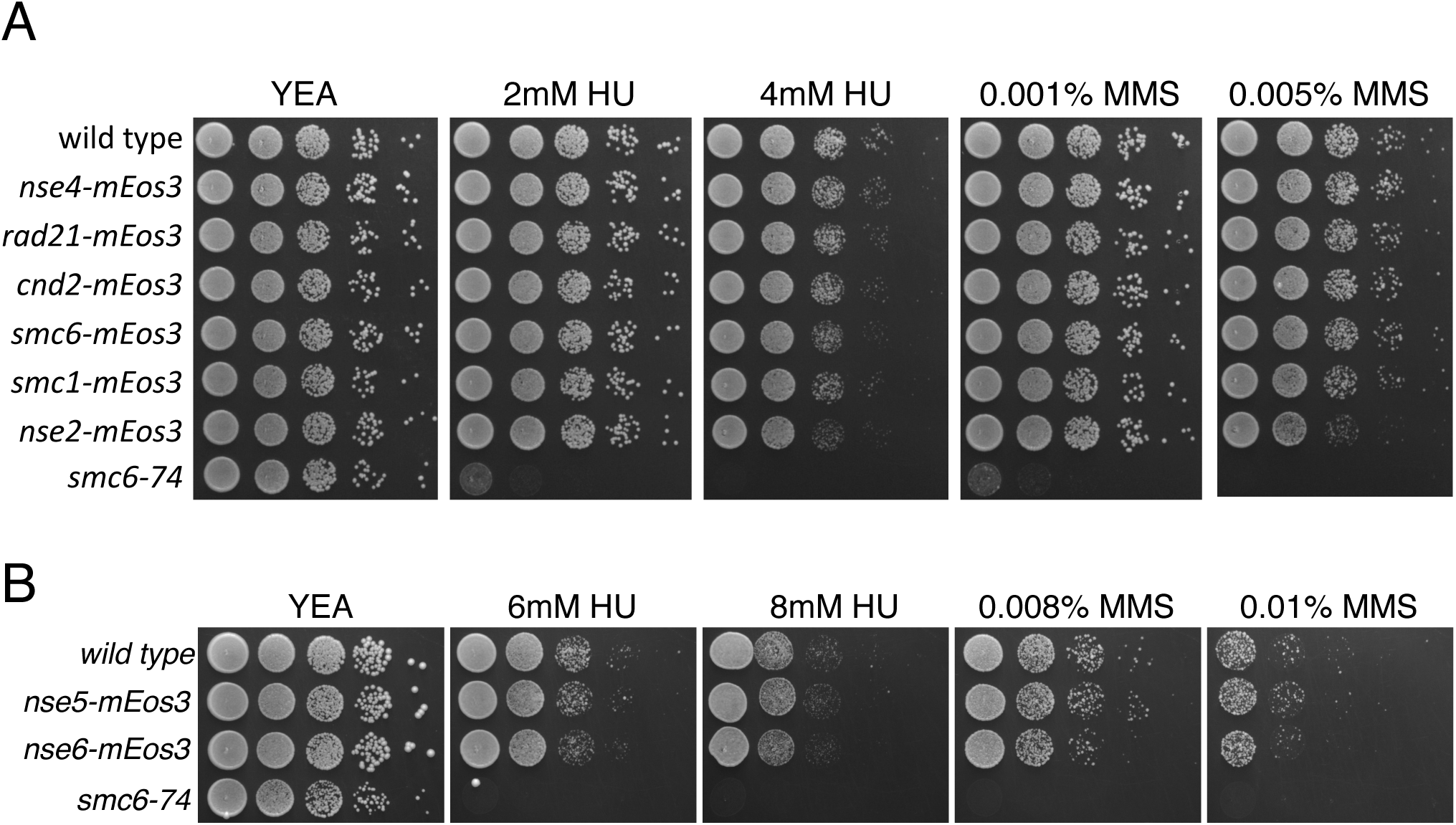
Characterisation of mEos3 tagged SMC subunits. Spot assay of *S. pombe* strains expressing A) different SMC components and B) Nse5 or Nse6 fused to the mEos3 fluorescent tag. Plates were incubated at 30°C for 3 days.

**Supplementary Figure 2.**
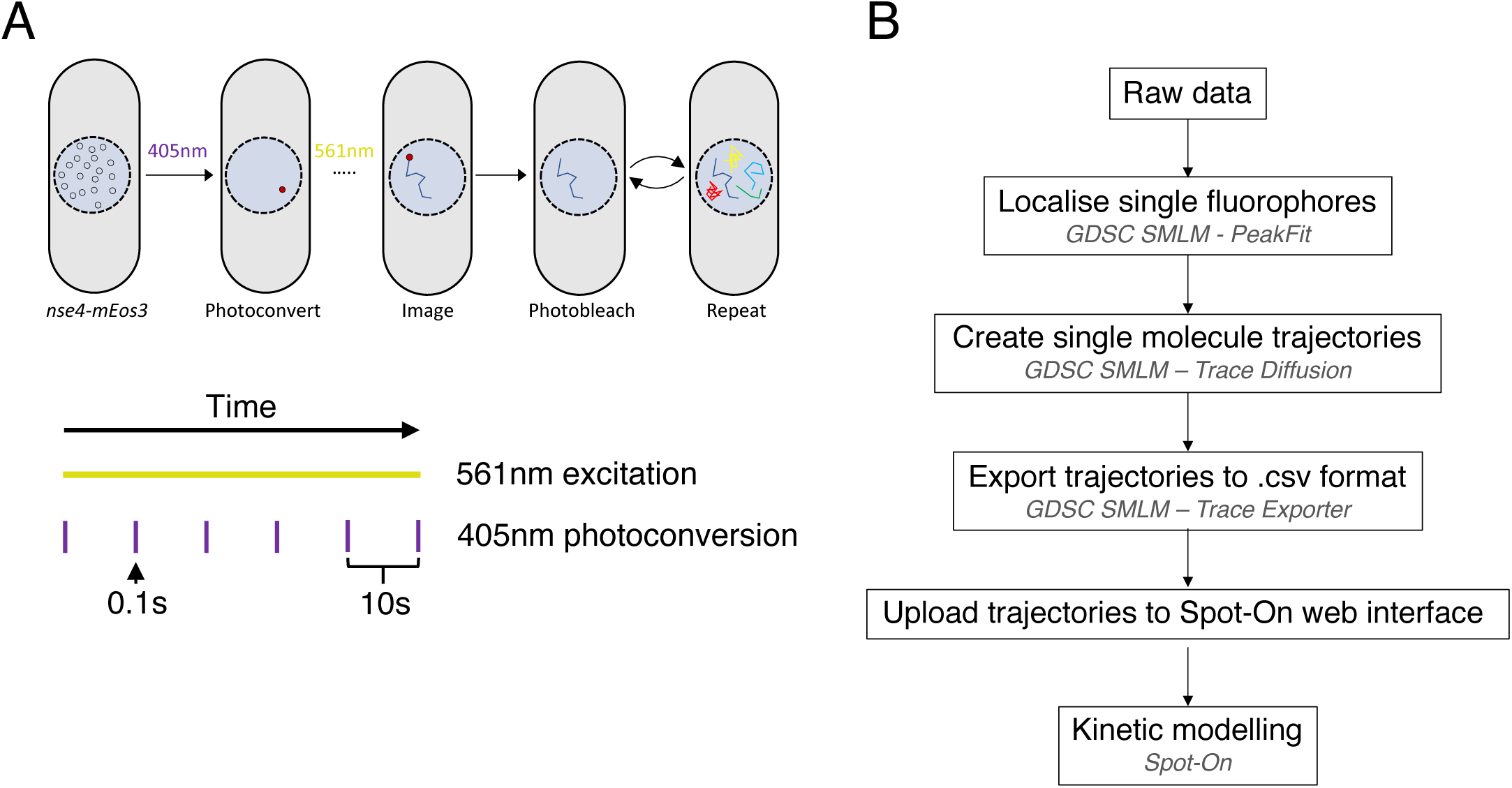
Outline of the single particle tracking technique. A. Diagram of SPT experimental approach. *Top* - Single mEos3 fluorophores fused to SMC components are stochastically photoconverted and imaged using 405nm and 561nm laser light respectively. In each frame the position of the fluorophore is recorded in each frame it is detected allowing for the creation of a trajectory. Multiple molecules are imaged in each cell over the course of an experiment. *Bottom* – Laser illumination scheme for each experiment. mEos3 is photoconverted using 0.1s pulses of 405nm laser every 10s and photoconverted species are imaged by continuous 561nm illumination. B. Raw data processing pipeline for SPT experiments. For specific details see Material and Methods.

**Supplementary Figure 3.**
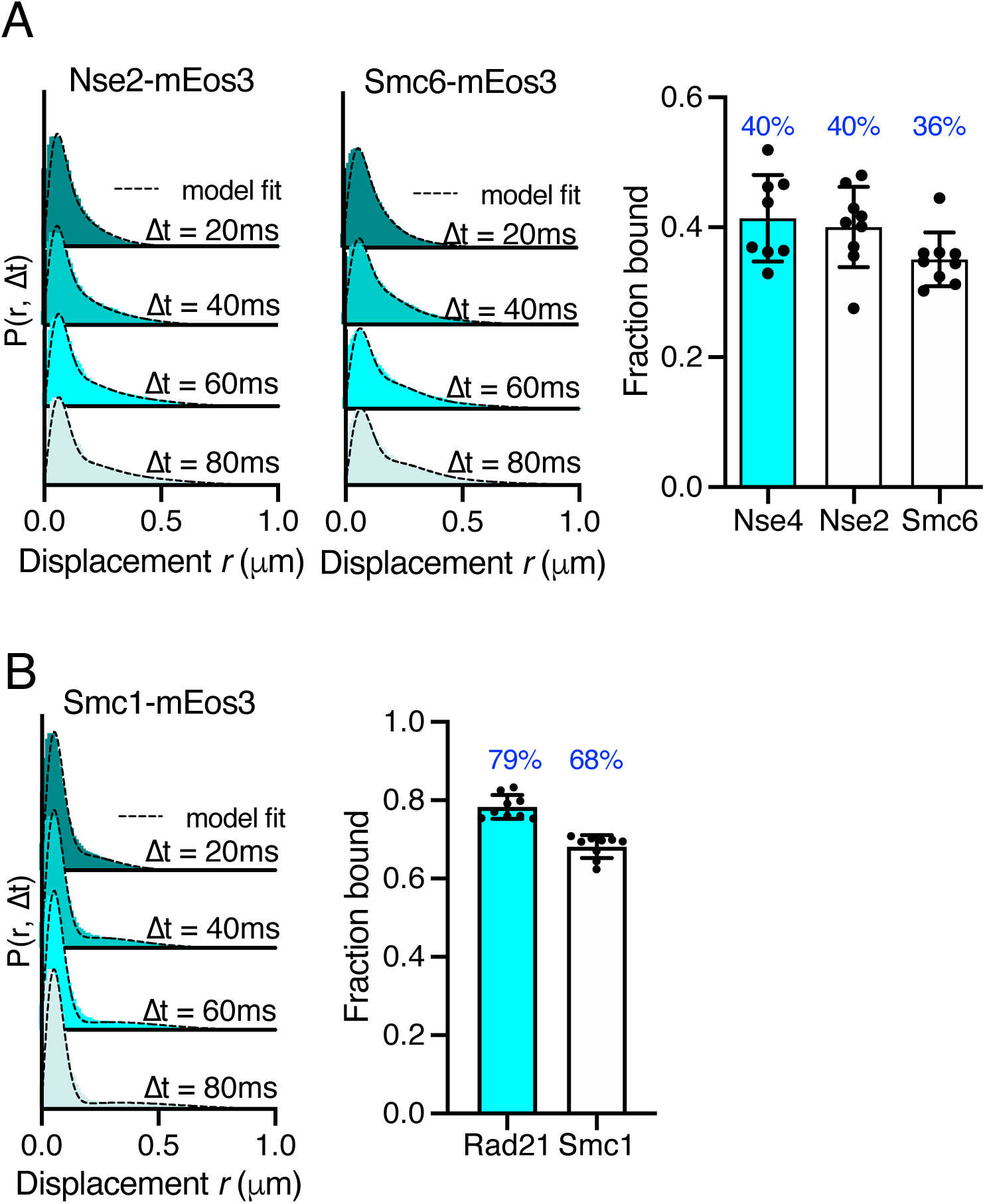
Single-particle tracking of SMC complex subunits. A. PDF histograms of single-molecule displacements (left) and fraction bound values calculated from Spot-On model fitting (right) of alternative Smc5/6 subunits tagged with mEos3 compared to Nse4-mEos3. Mean (+/− S.D). Black dots values derived from independent technical repeats, percentages in blue denote fraction bound value from fitting pooled data from all repeats B. PDF histograms of single-molecule displacements (left) and fraction bound values calculated from Spot-On model fitting (right) of alternative cohesin subunit, Smc1, tagged with mEos3 compared to Rad21-mEos3. Mean (+/− S.D). Black dots values derived from independent technical repeats, percentages in blue denote fraction bound value from fitting pooled data from all repeats

**Supplementary Figure 4.**
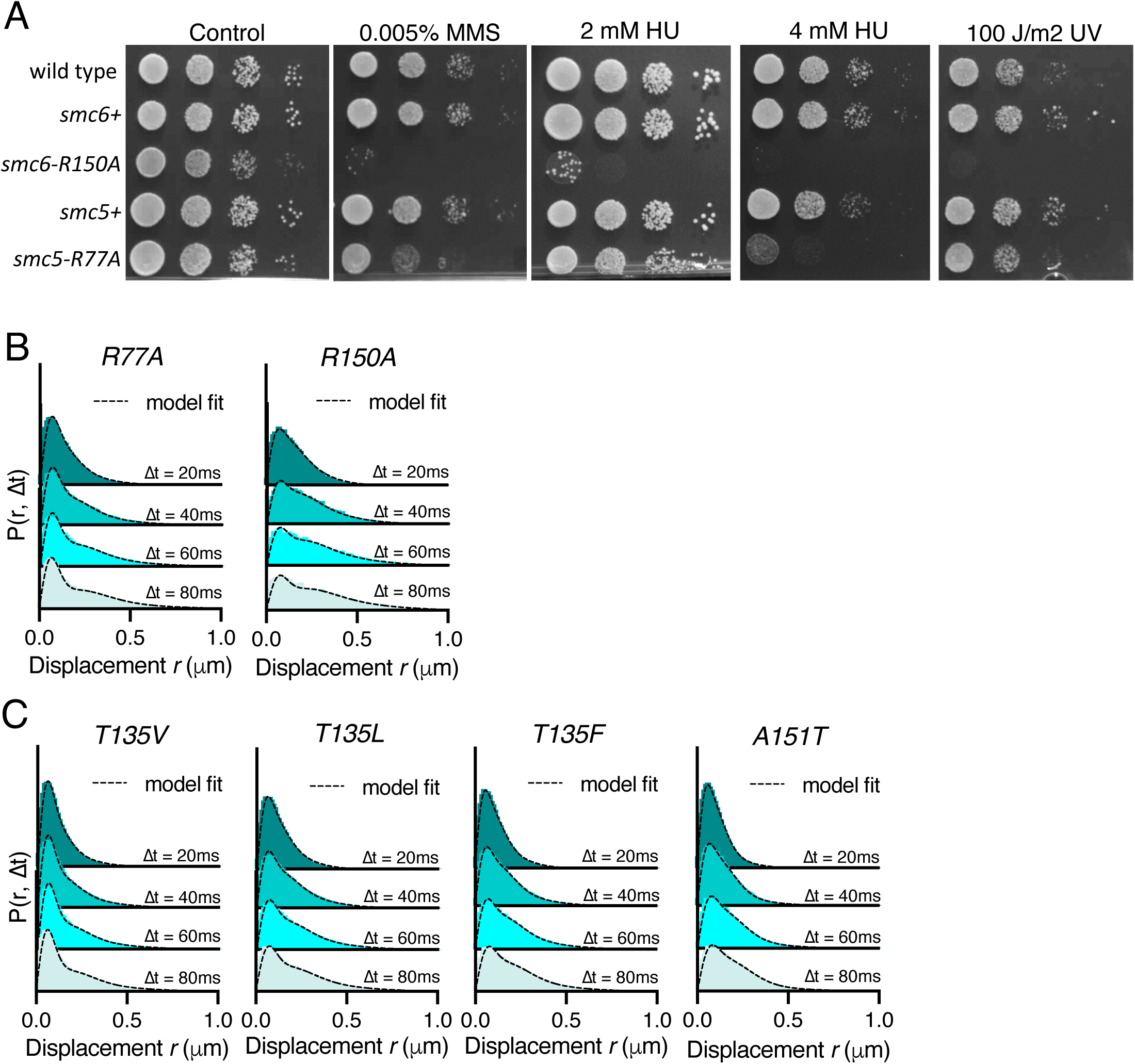
Characterisation of mEos3 tagged Smc5/6 ATPase mutants. A. Spot assay of *S. pombe* strains harbouring arginine finger mutations in either *smc5* or *smc6*. Plates were incubated at 30°C for 3 days. B. PDF histograms of single-molecule displacements for multiple Δt of Nse4-mEos3 for *smc5* (R77A) or *smc6* (R150A) arginine finger mutants (see Figure 3C and F for fraction bound). Dashed line indicates model derived from CDF fitting in Spot-On. Data are pooled from 3 individual experiments, each with 3 technical repeats. C. PDF histograms of single-molecule displacements for multiple Δt of Nse4-mEos3 in the indicated mutants (see figure 4E for fraction bound). Dashed line indicates model derived from CDF fitting in Spot-On. Data are pooled from 3 individual experiments, each with 3 technical repeats.

**Supplementary Figure 5.**
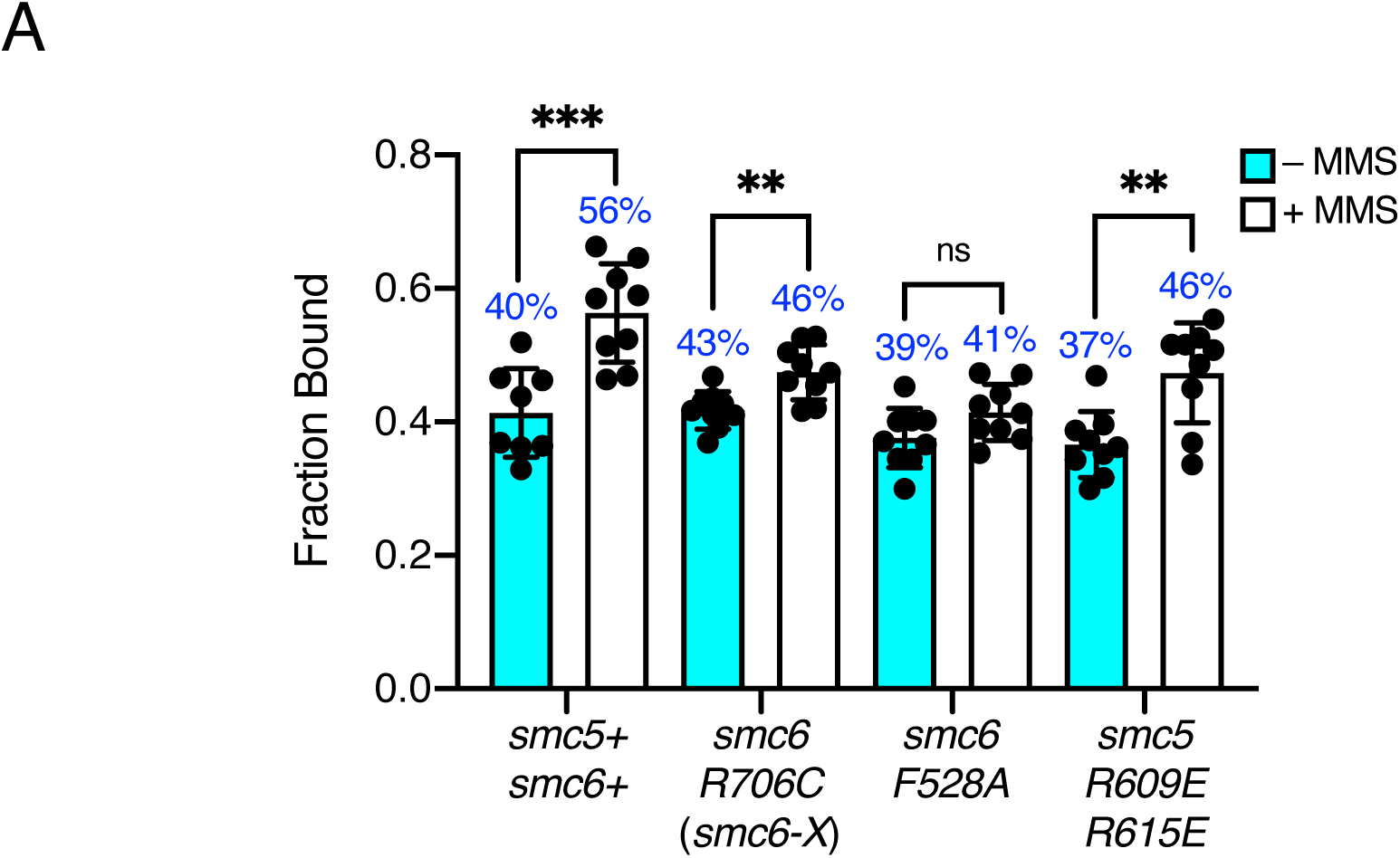
Spot-On analysis of MMS treated ssDNA interaction mutants. A. Fraction of bound Nse4-mEos3 extracted from Spot-On analysis of Nse4-mEos3 SPT in *smc6* hinge mutants treated with 0.03% MMS for 5 hours. Compared to asynchronous untreated datasets from Figure 4B. Mean +/− S.D. Black dots denote independent technical repeats, percentages denote fraction bound value from fitting pooled data from all repeats. ** = p< 0.005 (*smc6-X* = 0.0034, *smc5-RR* = 0.0025), *** p=0.0005.

**Supplementary Figure 6.**
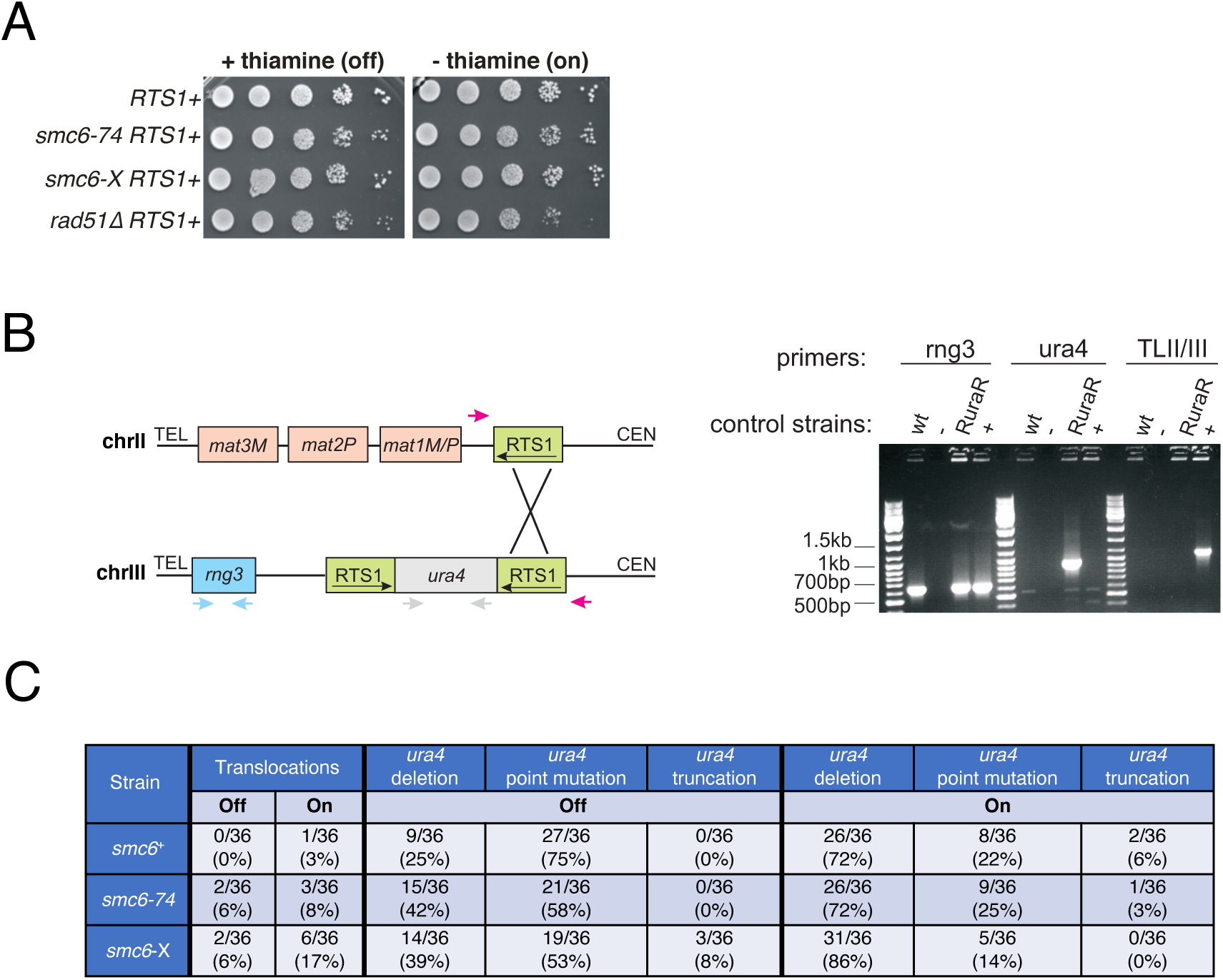
Analysis of the consequences of site-specific replication fork stalling on cell viability and gross chromosomal re-arrangements. **A.** Yeast spot assay of *S. pombe* strains harbouring the site-specific replication stall system *RuraR*. Replication fork stalling at *RTS1* is induced in the absence of thiamine (on). Plates were incubated at 30°C for 3 days. Unlike the HR-defective *rad51*Δ strain, *smc6* hypomorphs do not lose viability on induction of replication stalling at RTS1 **B.** PCR-based assay for translocation between *RTS1* at *RuraR* and the native *RTS1* at the mating type locus in *ura4^-^* colonies generated in the *ura4* loss of gene function assay^40^. Left: schematic to show the three primer pairs used. One pair (red arrows) amplifies the junction resulting from ectopic recombination between chromosome II and III (TLII/III). The second pair (grey arrows) amplifies the *ura4* locus to distinguish point mutations, truncations (internal deletions) and full-length deletions. *rng3* (blue arrows), an essential gene located between *RuraR* and the telomere, is amplified as positive control. Right: Example of control PCRs (top) and PCRs of 5-FOA resistant/*ura4^-^* colonies (bottom). The *rng3* product is amplified in all strains, but not in the negative control (“-”). *ura4* is amplified only in a *RuraR* strain, but not in Wild type (wt) (harbours full deletion of *ura4*, *ura4-D18*), the translocation positive control (“+”, gift from S. Lambert^40^) or the negative control. Translocation between chromosome II and III can only be detected in the positive control. **C.** PCR assay results for *ura^-^* colonies of *smc6+, smc6-*74 and *smc6-*X derived from the *RuraR ura4* loss assay carried out in the presence (*RuraR* arrest ‘Off’) or absence (*RuraR* arrest ‘On’) of thiamine.

**Supplementary Figure 7.**
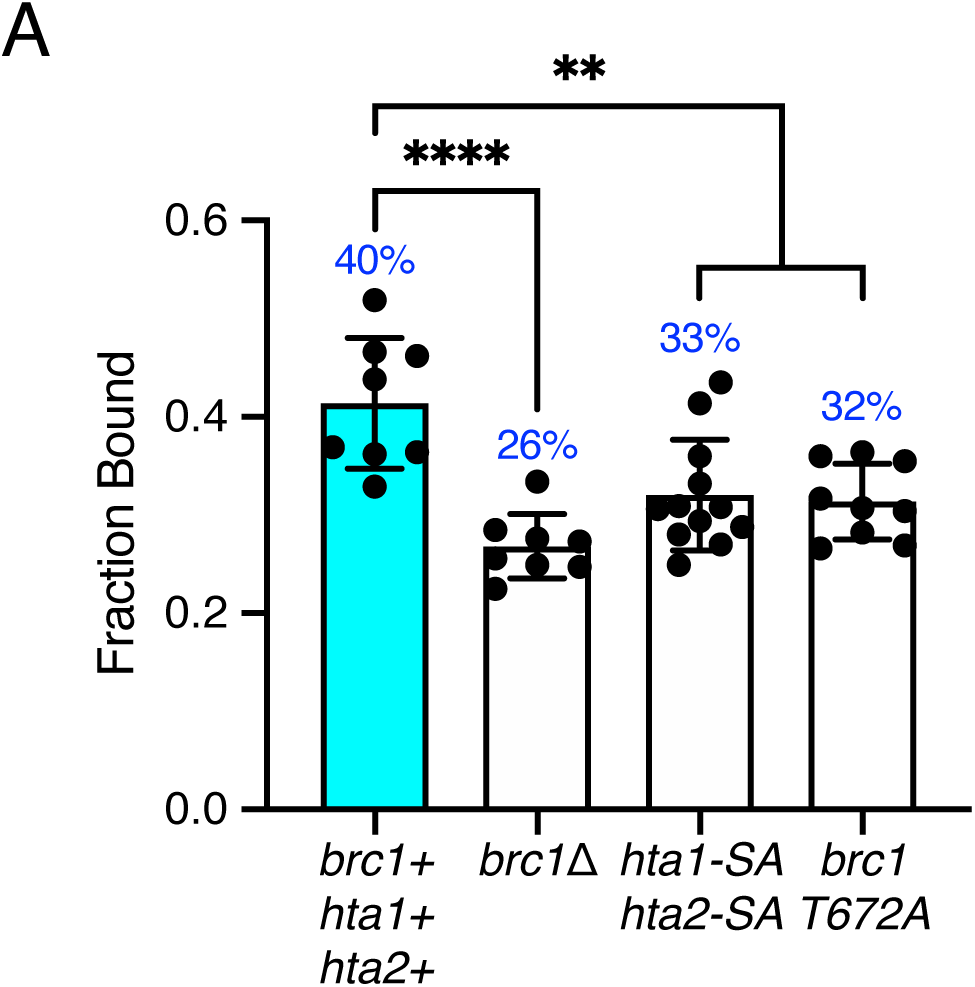
Spot-On analysis of Nse4 chromatin association in histone phosphorylation site and brc1γ-H2A interaction mutants. A. Fraction of bound Nse4-mEos3 extracted from Spot-On analysis of Nse4-mEos3 SPT in *hta1-S128A hta2-S129A* and *brc1-T672A* mutants compared to wild type and *brc1*Δ data sets from Figure 5C. Mean +/− S.D. Black dots denote independent technical repeats, percentages denote fraction bound value from fitting pooled data from all repeats. ** = p< 0.005 (*hta1-SA hta2-SA* = 0.0034, *smc5-RR* = 0.0016), **** p<0.0001.

**Supplementary Figure 8.**
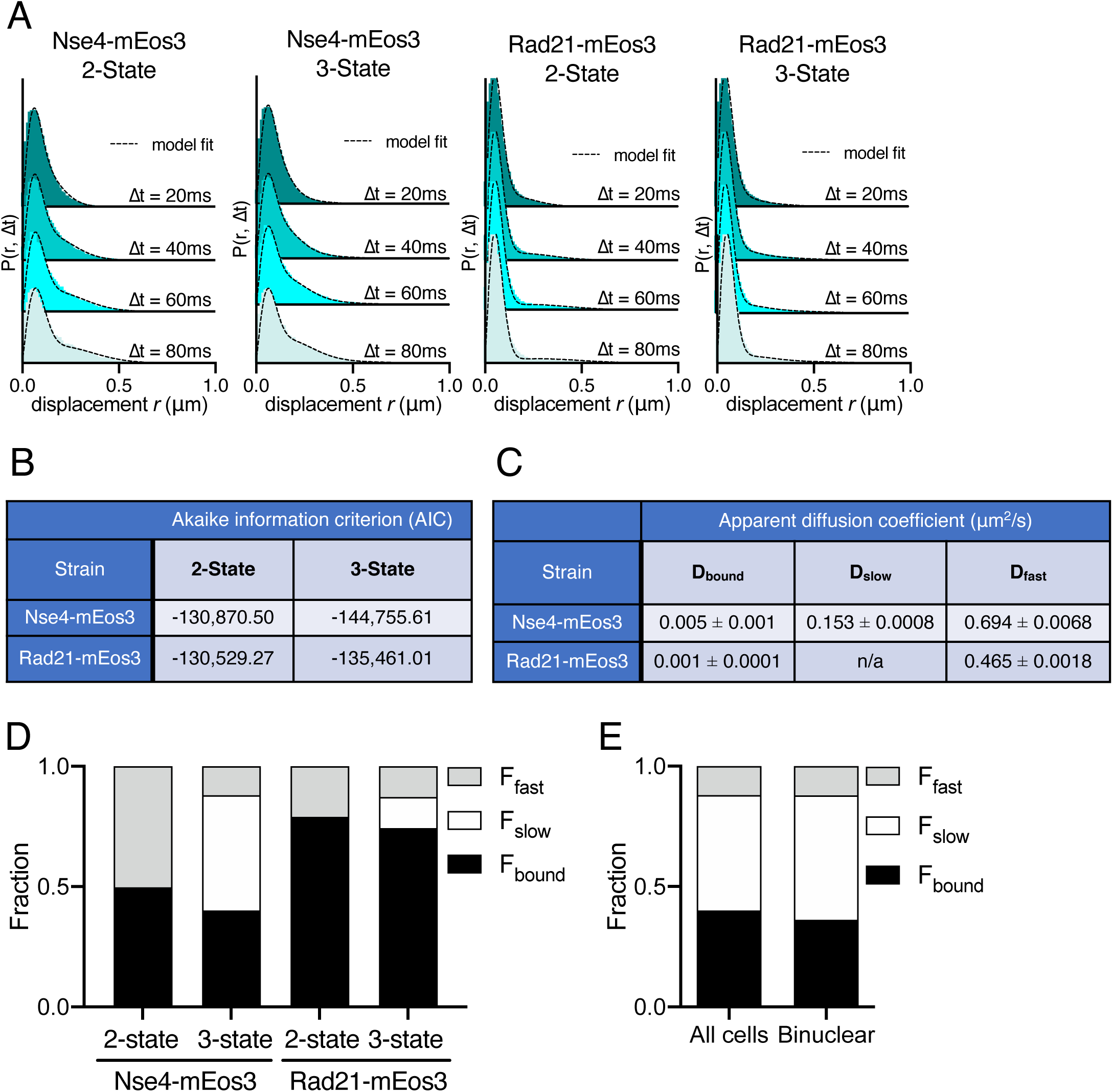
Smc5/6 behaviour fits a 3-state model. A. PDF histograms of single-molecule displacements for Nse4-mEos3 and Rad21-mEos3 over multiple Δt fit with either a 2-state or 3-state Spot-On model. Data are pooled from 3 independent experiments, each with three technical repeats. Dashed line indicates model derived from CDF fitting in Spot-On. B. Akaike information criterion (AIC) scores from Spot-On model fitting in A. Nse4-mEos3 3-state fitting showed a large difference in AIC scores compared to 2-state fitting. This indicates the data are best described by a 3-state model. The difference in AIC scores for Rad21-mEos3 was much smaller and thus a 2-state model was used. C. Apparent diffusion coefficients of Spot-On sub-populations of Nse4-mEos3 (3-State) and Rad21-mEos3 (2-State). D. Fractions of the total population of molecules observed residing in each kinetic state extracted from Spot-On model fitting data in A. E. Comparison of the fractions of Nse4-mEos3 molecules observed residing in each kinetic state extracted from all cells (mostly G2) in the wild type data set or only binuclear cells (S-phase, n=75).

## Supplementary Tables

**Table S1.** Fluctuation experiment data. Data from individual experimental repeats of *ura4* loss assay in Figure 4C

| **Experiment 1** Rate of *ura4* loss per cell per generation |  |  |  |
| --- | --- | --- | --- |
| Strain | Off<br>(+thiamine) | On<br>(-thiamine) | Relative<br>to Off |
| *smc6*<sup>+</sup> | 2.2 x 10<sup>-8</sup> | 4.6 x 10<sup>-8</sup> | 2.09 |
| *smc6-74* (A151T) | 1.4 x 10<sup>-7</sup> | 3.83 x 10<sup>-7</sup> | 2.71 |
| *smc6-X* (R706C) | 8.4 x 10<sup>-9</sup> | 1.6 x 10<sup>-6</sup> | 194.00 |
| *nse3-R254E* | 6.0 x 10<sup>-8</sup> | 2.4 x 10<sup>-7</sup> | 3.99 |

| **Experiment 2** Rate of *ura4* loss per cell per generation |  |  |  |
| --- | --- | --- | --- |
| Strain | Off<br>(+thiamine) | On<br>(-thiamine) | Relative<br>to Off |
| *smc6*<sup>+</sup> | 3.2 x 10<sup>-8</sup> | 5.7 x 10<sup>-8</sup> | 1.77 |
| *smc6-74* (A151T) | 2.3 x 10<sup>-7</sup> | 1.56 x 10<sup>-6</sup> | 6.78 |
| *smc6-X* (R706C) | 1.3 x 10<sup>-7</sup> | 7.7 x 10<sup>-6</sup> | 61.2 |
| *nse3-R254E* | 7.2 x 10<sup>-8</sup> | 6.0 x 10<sup>-7</sup> | 8.36 |

| **Experiment 3** Rate of *ura4* loss per cell per generation |  |  |  |
| --- | --- | --- | --- |
| Strain | Off<br>(+thiamine) | On<br>(-thiamine) | Relative<br>to Off |
| *smc6*<sup>+</sup> | 1.6 x 10<sup>-8</sup> | 1.8 x 10<sup>-8</sup> | 1.12 |
| *smc6-74* (A151T) | 7.09 x 10<sup>-8</sup> | 3.98 x 10<sup>-7</sup> | 5.61 |
| *smc6-X* (R706C) | 1.3 x 10<sup>-7</sup> | 1.4 x 10<sup>-5</sup> | 105.70 |
| *nse3-R254E* | 1.3 x 10<sup>-8</sup> | 4.1 x 10<sup>-7</sup> | 32.20 |

**Table S2.** Strain table. Strains used during this study

| Strain No. | Genotype | Reference |
| --- | --- | --- |
| TJE323 | *loxP-nse4-mEos3.2-loxM3 ura4-D18 leu1-32 ade6-704* | This study |
| TJE350 | *loxP-smc6-mEos3.2-loxM3 ura4-D18 leu1-32 ade6-704* | This study |
| TJE496 | *smc1-loxP-mEos3.2:kanMX6-loxM3 ura4-D18 leu1-32* | This study |
| TJE480 | *loxP-nse4-mEos3.2-loxM3 loxP-smc6-T135V-loxM3 ura4-D18 leu1-32 ade6-704* | This study |
| TJE477 | *loxP-nse4-mEos3.2-loxM3 loxP-smc6-T135L-loxM3 ura4-D18 leu1-32 ade6-704* | This study |
| TJE475 | *loxP-nse4-mEos3.2-loxM3 loxP-smc6-T135F-loxM3 ura4-D18 leu1-32 ade6-704* | This study |
| TJE410 | *loxP-nse4-mEos3.2-loxM3 smc6-A151T ura4-D18 leu1-32 ade6-704* | This study |
| TJE719 | *loxP-nse4-mEos3.2-loxM3 loxP-smc6-R150A-loxM3 ura4-D18 leu1-32 ade6-704* | This study |
| TJE711 | *loxP-nse4-mEos3.2-loxM3 loxP-smc5-R77A-loxM3 ura4-D18 leu1-32 ade6-704* | This study |
| TJE509 | *loxP-nse4-mEos3.2-loxM3 smc6-F528A ura4-D18 leu1-32 ade6-704* | This study |
| TJE483 | *loxP-nse4-mEos3.2-loxM3 smc5-R609E R615E ura4-D18 leu1-32 ade6-704* | This study |
| TJE671 | *loxP-nse4-mEos3.2-loxM3 smc5-Y612G ura4-D18 leu1-32 ade6-704* | This study |
| TJE418 | *loxP-nse4-mEos3.2-loxM3 smc6-R706C ura4-D18 leu1-32 ade6-704* | This study |
| TJE492 | *loxP-nse4-mEos3.2-loxM3 loxP-nse3-R254E-loxM3 ura4-D18 leu1-32 ade6-704* | This study |
| TJE730 | *loxP-nse4-mEos3.2-loxM3 brc1::hphMX6 ura4-D18 leu1-32 ade6-704* | This study |
| TJE734 | *loxP-nse4-mEos3.2-loxM3 nse6::kanMX6 ura4-D18 leu1-32 ade6-704* | This study |
| TJE796 | *nse6-loxP-mEos3.2-loxM3 ura4-D18 leu1-32* | This study |
| TJE816 | *loxP-nse4-mEos3.2-loxM3 hta1-S129A:ura4 hta2-S128A:his3 his3-D1 ura4-D18 leu1-32* | This study |
| TJE393 | *rad21-loxP-mEos3.2:kanMX6-loxM3 ura4-D18 leu1-32* | This study |
| TJE586 | *nse2-loxP-mEos3.2:kanMX6-loxM3 ura4-D18 leu1-32* | This study |
| TJE522 | *cnd2-loxP-mEos3.2:kanMX6-loxM3 ura4-D18 leu1-32* | This study |
| TJE886 | *loxP-nse4-mEos3.2-loxM3 mcm4-loxP-yEGFP:KanMX6-loxM3 ura4-D18 leu1-32 ade6-704* | This study |
| THE 884 | *loxP-nse4-mEos3.2-loxM3 loxP-brc1-T672A-loxM3 ura4-D18 leu1-32* | This study |
| TJE828 | *nse6-loxP-mEos3.2-loxM3 smc6-74 ura4-D18 leu1-32* | This study |
| TJE830 | *nse6-loxP-mEos3.2-loxM3 smc6-X ura4-D18 leu1-32* | This study |
| TJE888 | *loxP-nse4-mEos3.2-loxM3 brc1::hphMX6 nse6::kanMX6 ura4-D18 leu1-32 ade6-704* | This study |
| HQD87 | *loxP-smc5+-ura4-loxM3 ura4-D18 leu1-32 ade6-704* | This study |
| DE297 | *loxP-smc6+-ura4-loxM3 ura4-D18 leu1-32 ade6-704* | This study |
| DE285 | *loxP-smc6-T135V-loxM3 ura4-D18 leu1-32 ade6-704* | This study |
| DE283 | *loxP-smc6-T135L-loxM3 ura4-D18 leu1-32 ade6-704* | This study |
| DE281 | *loxP-smc6-T135F-loxM3 ura4-D18 leu1-32 ade6-704* | This study |
| DE279 | *loxP-smc6-R150A-loxM3 ura4-D18 leu1-32 ade6-704* | This study |
| DE342 | *loxP-smc5-R77A-loxM3 ura4-D18 leu1-32 ade6-704* | This study |
| DE273 | *smc6-A151T ura4-D18 leu1-32 ade6-704* | Lab strain |
| JMM1188 | *ura4-D18 leu1-32 ade6-704* | Lab strain |
| JMM1162 | *nmt41:rtf1:sup35 RTS1-ura4-RTS1 ade6-704 leu1-32* | Lambert et al 2005 |
| JMM1171 | *rhp51::NAT nmt41:rtf1:sup35 RTS1-ura4-RTS1 ade6-704 leu1-32* | Lambert et al 2005 |
| JMM1371 | *smc6-A151T nmt41:rtf1:sup35 RTS1-ura4-RTS1 ade6-704 leu1-32* | This study |
| JMM1375 | *smc6-R706C nmt41:rtf1:sup35 RTS1-ura4-RTS1 ade6-704 leu1-32* | This study |
| DE331 | *nse3-R254E nmt41:rtf1:sup35 RTS1-ura4-RTS1 ade6-704 leu1-32* | This study |

